# ARF6-dependent endocytic trafficking of the Interferon-γ receptor drives adaptive immune resistance in cancer

**DOI:** 10.1101/2023.09.29.560199

**Authors:** Yinshen Wee, Junhua Wang, Emily C. Wilson, Coulson P. Rich, Aaron Rogers, Zongzhong Tong, Evelyn DeGroot, Y.N. Vashisht Gopal, Michael A. Davies, H. Atakan Ekiz, Joshua K.H. Tay, Chris Stubben, Kenneth M. Boucher, Juan M. Oviedo, Keke C. Fairfax, Matthew A. Williams, Sheri L. Holmen, Roger K. Wolff, Allie H. Grossmann

## Abstract

Adaptive immune resistance (AIR) is a protective process used by cancer to escape elimination by CD8^+^ T cells. Inhibition of immune checkpoints PD-1 and CTLA-4 specifically target Interferon-gamma (IFNγ)-driven AIR. AIR begins at the plasma membrane where tumor cell-intrinsic cytokine signaling is initiated. Thus, plasma membrane remodeling by endomembrane trafficking could regulate AIR. Herein we report that the trafficking protein ADP-Ribosylation Factor 6 (ARF6) is critical for IFNγ-driven AIR. ARF6 prevents transport of the receptor to the lysosome, augmenting IFNγR expression, tumor intrinsic IFNγ signaling and downstream expression of immunosuppressive genes. In murine melanoma, loss of ARF6 causes resistance to immune checkpoint blockade (ICB). Likewise, low expression of ARF6 in patient tumors correlates with inferior outcomes with ICB. Our data provide new mechanistic insights into tumor immune escape, defined by ARF6-dependent AIR, and support that ARF6-dependent endomembrane trafficking of the IFNγ receptor influences outcomes of ICB.

## Introduction

Immune escape, a hallmark of cancer ^1^, includes cancer cell sensing and directly disarming immune attack. Tumors can adapt to immune editing by expressing and activating immunosuppressive molecules ^2,3^. For example, Interferon gamma (IFNγ), secreted by cytotoxic T cells and natural killer cells, activates anticancer immunity in the tumor microenvironment and incites tumor-intrinsic JAK-STAT signaling, which in the acute setting can have cytostatic and cytotoxic effects on cancer cells ^4^. Chronic exposure to IFNγ elicits adaptive expression of immunosuppressive genes from cancer cells, including *CD274* (encoding PD-L1), *CD80* and *IDO1* ^3,5,6^. PD-L1 and CD80 are immune checkpoint ligands for PD-1 and CTLA-4 receptors, respectively, expressed on cytotoxic CD8^+^ T cells to inhibit their activity. Immune checkpoint blockade (ICB) therapies pharmacologically block binding of PD-L1 and CD80 to their respective receptors and disrupt these pathways, alleviating immune suppression and restoring the effector function of cytotoxic T cells. IFNγ-driven AIR is a broadly conserved mechanism of immune escape in cancer, highlighted by the clinical utility of ICB in numerous cancer types. In addition to cutaneous melanoma, ICB has been approved to treat several types of carcinomas, mesothelioma, a subset of hematopoietic malignancies, and sarcomas ^7,8^. Unfortunately, only about 25% of patients with advanced solid tumors treated with ICB respond ^3^. Mechanisms that drive low response rates remain incompletely understood.

AIR begins at the plasma membrane of cancer cells when they are exposed to IFNγ, Interferon alpha (IFNα), Tumor Necrosis Factor alpha (TNFα), lytic granules and death receptor ligands from cytotoxic T lymphocytes (CTLs) ^2^. The plasma membrane is a dynamic interface where a repertoire of proteins and lipids is remodeled by the endomembrane trafficking system in response to changing environments and cellular needs ^9^. Endomembrane trafficking machineries are responsible for internalizing plasma membrane proteins and directing them towards being secreted, recycled to the cell surface, or degraded in lysosomes. Given the rush of cytokines released during immune attack, cancer cells might dynamically modulate cytokine receptor density at their surface in an attempt to adapt and survive. At present, our understanding of this process and how it impacts tumor progression and response to ICB is limited.

The small GTPase ADP-Ribosylation Factor 6 (ARF6) localizes to the plasma membrane where it mediates endocytosis and recycling of plasma membrane proteins ^10–13^. ARF6 is activated by and coordinates signaling, cargo transport and functional output of diverse ligand-receptor systems ^14–21^. In addition, ARF6 is upregulated and/or activated downstream of oncoproteins such as mutant GNAQ ^22^, p53 and KRAS ^23^. Like other small GTPases, activation of ARF6 by GTP loading is mediated by guanine exchange factors (GEFs), while conversion of active ARF6-GTP to inactive ARF6-GDP is catalyzed by GTPase activating proteins (GAPs) (Figure 1A). Thus, GEFs and GAPs determine the lifetime of ARF6 activation and an imbalance in expression of these proteins could shift the activation-deactivation cycle of ARF6 to favor one state over the other. In fact, we reported that reduced expression of ARF6 GAPs (*ACAP1 and ARAP2*), in metastatic melanoma from Stage III patients, was associated with inferior overall survival^24^. We also showed that ARF6-GTP levels were aberrantly high in metastatic melanoma, compared to adjacent benign tissues, and that ARF6 activation accelerated spontaneous metastasis in xenografts and genetically engineered tumor models^15,24^. Specifically, ARF6-GTP in primary tumors promoted metastasis without altering primary tumor growth. This result can be partly attributed to the pro-invasive functions of ARF6 that we and others have reported^15,24^, however, this may not fully explain the pro-metastatic roles of ARF6. Successful metastasis requires much more than the acquisition of invasive behavior; it also requires immune escape. Hence, ARF6-mediated endocytic trafficking of cytokine receptors, or other immunomodulating surface proteins, could be involved in immune escape during primary tumor development, facilitating metastasis.

**Figure 1:**
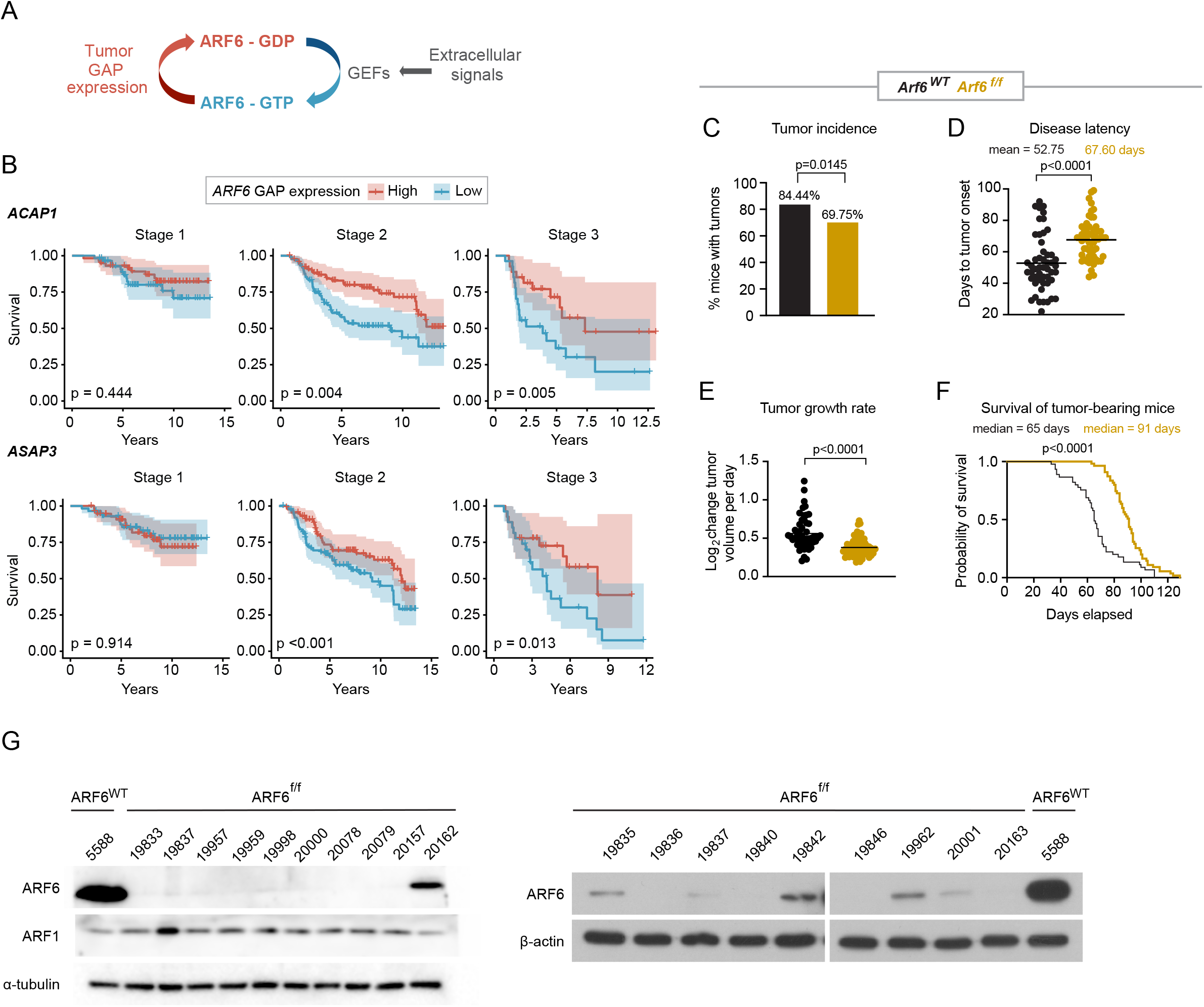
ARF6 promotes primary melanoma formation and progression. **(A)** Schematic diagram showing ARF6 cycles between the GDP-bound inactive form (red) and the GTP-bound active form (blue). **(B)** Correlations between the top and the bottom quartile of mRNA expression levels of indicated ARF6 GAPs in primary cutaneous melanoma (the Leeds cohort) with survival, n= 350 patients. Cox proportional hazards regression. **(C-F)** Melanocyte-specific deletion of *Arf6* restricts tumorigenesis. **(C)** Percent of *Dct-TVA; Braf^V^*^600^*^E^; Cdkn2a^f/f^* mice that developed tumors within 100 days after Cre injection (tumor induction). n=90 *Arf6* wild type (*Arf6^WT^*), n= 119 *Arf6* floxed (*Arf6^f/f^*), two-sided Fisher’s exact test. **(D)** Days to initial tumor detection after Cre injection, two-tailed t-test with Welch’s correction. **(E)** Rate of tumor growth measured from time of initial detection, **(F)** Survival of mice (before primary tumor reached 2cm) after Cre injection (day 0) within 130 days, n=45 controls, n=54 *Arf6^f/f^*, Log-rank (Mantle-Cox) test. **(G)** Western blot detection of indicated proteins in early-passage primary tumor cell lines. See also Figure S1A.

Melanoma has a high propensity for early metastasis, when primary tumors are as thin as one millimeter^25^, indicating that the behavior of melanoma cells in early development is tightly linked to metastatic progression. Cumulative evidence demonstrates that this behavior at least partly results from an innate ability of malignant melanocytes to suppress and evade the immune system. For example, PD-L1 expression in primary cutaneous melanomas is associated with CD8^+^ T cell tumor infiltration^26^ and in high-risk Stage II (non-metastatic) melanoma patients, adjuvant anti-PD-1 therapy significantly improves recurrence free survival and distant metastasis free survival^27^. These data suggest that IFNγ-mediated AIR can drive progression of early-stage disease. More recently, Nirmal et al.^28^ reported that CTLs, regulatory T cells (Tregs) and myeloid cells were detectable in atypical (precursor) melanocytic lesions and increased with progression to melanoma *in situ* (antecedent to invasion). At the invasive stage, the tumor-immune interface had evolved into complex geospatial microenvironments where multiple mechanisms of immune evasion prevailed. These data provide evidence that newly transformed melanocytes are subject to immune editing and have an intrinsic ability to alter the microenvironment to favor an immunosuppressive state, a process that is poorly understood.

To better understand how ARF6 shapes the development and progression of cancer in an immunocompetent host, we expanded our genetically engineered murine models^24^ to create BRAF^V600E^ melanoma with melanocyte-specific, inducible deletion of the *Arf6* gene. Using these models together with human tumor cells, we report here that ARF6 is activated by IFNγ and is necessary for trafficking of the IFNγ receptor (IFNγR), controlling receptor protein levels in melanoma and other cancer lineages. As a result, ARF6 shapes IFNγ-driven AIR during tumor progression, which in turn determines the response to ICB.

## Results

### ARF6 promotes primary melanoma formation and progression

Given our previous data, we asked whether ARF6 might be aberrantly activated and play a critical role during primary tumor progression. Similar to our previous analysis of TCGA metastatic melanomas ^24^, we interrogated expression of *ARF* as well as ARF GEF and GAP genes (Table S1) in primary tumors from the Leeds Melanoma Cohort, which includes over 700 primary melanomas from Stage I-III, treatment naïve patients ^29^. For unknown reasons, *ARF6* was not included in the tumor gene expression dataset. Nevertheless, similar to metastatic melanoma, low expression of ARF6 GAP genes in these primary tumors significantly correlated with reduced overall survival of Stage II and Stage III patients (Figure 1B). The prognostic ARF GAP genes include *ACAP1* ^30^, an ARF6 GAP, and *ASAP3*, an ARF1/ARF5/ARF6 GAP^31^. Interestingly, *ACAP1* is prognostic in both primary (Leeds cohort, Figure 1B) and metastatic (TCGA cohort)^24^ tumors.

To investigate a role for ARF6 in the development and progression of primary melanoma *in vivo*, we crossed *Arf6^flox/flox^* (*Arf6^f/f^*) mice^32^ with *Dct:TVA; Braf^V^*^600^*^E^;Cdkn2a^f/f^* mice and induced genetic alterations specifically in melanocytes via subcutaneous injection of RCAS-Cre into the flank, as previously described ^33^. In this model, loss of *Arf6* significantly reduced tumor incidence (Figure 1C), increased disease latency (Figure 1D), slowed tumor growth, which was measured from the time of tumor formation (Figure 1E), and prolonged host survival (Figure 1F). Interestingly, Western blot of primary tumor cell lines showed that up to 30% of the *Arf6*^f/f^ tumors retained a comparatively low level of ARF6 expression, whereas ARF1 expression remained intact (Figure 1G). These data suggest that in a fraction of the *Arf6^f/f^* mice, *Arf6^WT^* tumor subclones may exist due to incomplete Cre-mediated recombination of the *Arf6* allele. Consistently, *in situ* hybridization detected reduced, heterogeneous (absent or multi focal) *Arf6* mRNA signals in whole tumor sections (tumor + intact stroma) from *Arf6^f/f^* mice (Figure S1A). Thus, low expression of *Arf6* may have persisted in a minor fraction of the *Arf6^f/f^* cohort, yet despite this, there is a significant defect in tumor development and growth in this population (Figures 1C-F). Interestingly, tumor-specific expression of ARF6^Q67L^, a constitutively active (GTP-bound) form of ARF6, accelerated spontaneous metastasis without altering primary tumor development, proliferation, or growth in this model ^24^. These combined data may reflect distinct functions for ARF6 that depend on expression level and/or activation state.

### Loss of ARF6 enhances tumor inflammation and apoptosis

To explore potential mechanisms of ARF6-dependent tumor progression, we analyzed pathway alterations using bulk transcriptomes from murine tumors expressing constitutively active ARF6^Q67L^ (phenotype previously published ^24^) or deleted *Arf6* (ARF6^f/f^), each compared to wild type ARF6 (ARF6^WT^) tumors. Strikingly, ARF6^Q67L^ and ARF6^f/f^ tumors shared several Hallmark gene sets, with opposite directions of enrichment (Figure S1B-S1C), highlighted by significantly decreased expression of IFNα, IFNγ and TNFα signatures in ARF6^Q67L^ tumors but enrichment of these in ARF6^f/f^ tumors (Figure 2A). These cytokine signatures suggest that ARF6 may control the ability of tumors to shape their immune microenvironment. Histologically, tumor infiltrating lymphocytes (TIL) were scattered diffusely in the ARF6^WT^ tumors and formed subtle, small clusters rarely; in contrast, the TIL in ARF6^f/f^ tumors formed obvious, robust clusters, and were evident in significantly more mice (Figure 2B). In addition, compared to ARF6^WT^ tumors, ARF6^f/f^ tumors showed significantly decreased levels of phosphorylation of death-associated protein kinase 1 (DAPK1) at serine 308, which is reported to initiate IFNγ-induced apoptosis^34^, and increased levels of cleaved executioner caspases 3 and 7 (Figure 2C), indicating increased apoptosis. Importantly, consistent with murine tumors, primary human melanomas with high expression of ARF6 GAPs, which are predicted to have relatively low ARF6-GTP levels, showed significant enrichment in tumor-infiltrating immune cells, particularly cytotoxic CD8^+^ T cells, compared to tumors with low expression of ARF6 GAPs (Figure 2D). These data together support that the overall level of ARF6 activation in tumor cells may have regulated an anti-tumor immune response.

**Figure 2:**
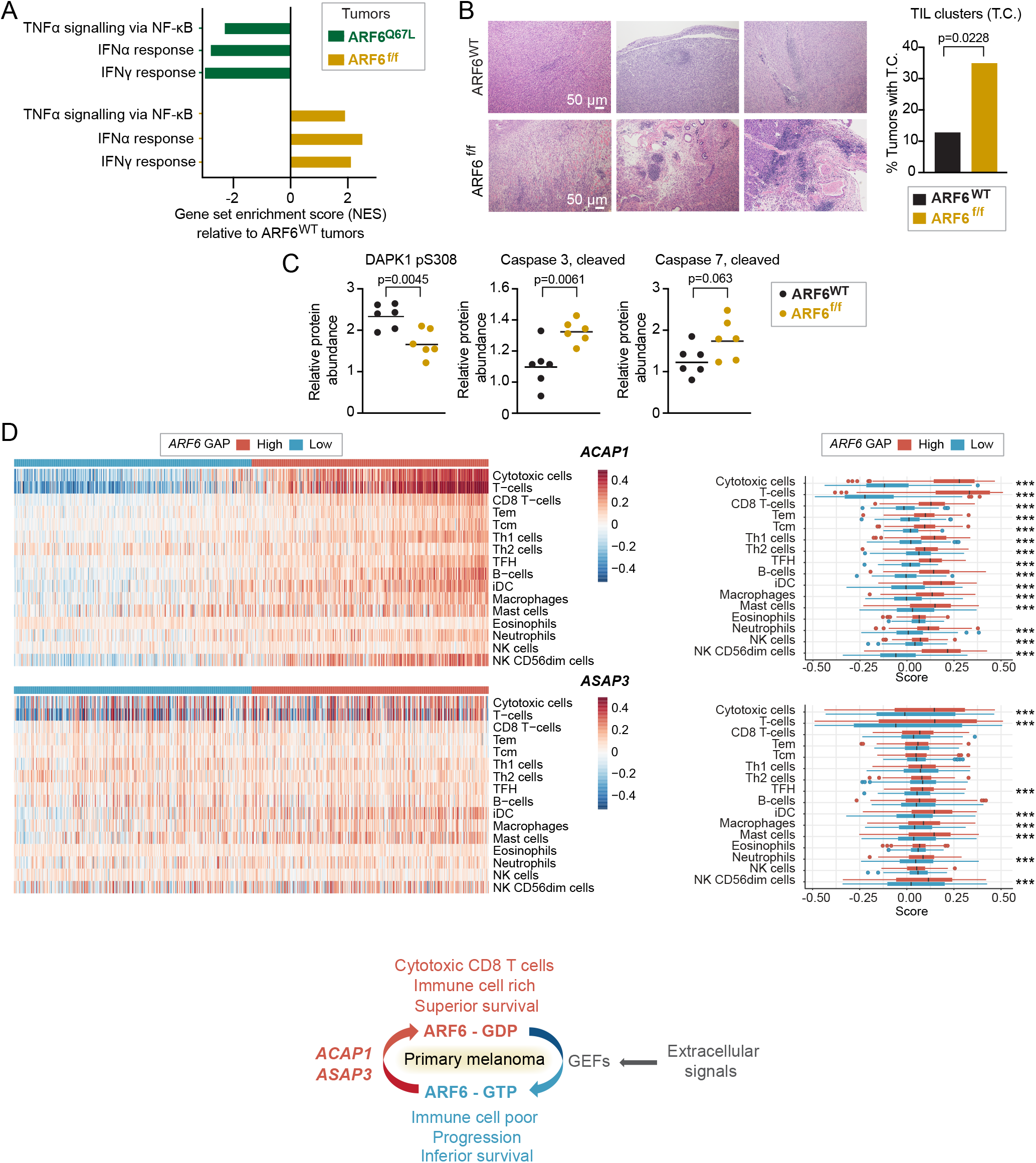
ARF6-dependent tumor inflammation and apoptosis. **(A)** Shared significantly enriched gene sets (MSigDB Hallmark), but in opposite directions, between ARF6^Q67L^ and ARF6^f/f^ tumors from bulk tumor transcriptomes (n=6 mice each). See also Figure S1B. **(B)** Representative images of H&E staining showing TIL clusters, scale bars = 50μm, and fractions of tumors with TIL clusters (n= 46 ARF6^WT^ controls, n=40 ARF6^f/f^, Fisher’s exact test, two-sided. **(C)** Apoptotic protein profile of tumors (n=6 mice each) detected by Reverse Phase Protein Array, two-tailed t-test. **(D)** Immune cell gene set enrichment in primary human melanoma (Leeds melanoma, n=350), supervised clustering with ARF6 GAP expression (related to Figure 1B). *** p<0.001. *ACAP1* cytotoxic T cells p=9.924×10^−124^, T cells p=2.636×10^−141^. *ASAP3* cytotoxic T cells p=2.7081×10^−8^, T cells p= 1.2997×10^−8^. Schematic = ARF6 activation cycle related to ARF6 GAP expression detected in primary tumors with associated immune cell signatures and survival outcome.

### ARF6 is critical for tumor intrinsic suppression of the adaptive immune response

To determine how ARF6 might alter the tumor immune microenvironment (TIME), we profiled the immune cells in ARF6^WT^ and ARF6^f/f^ tumors using flow cytometry. While the absolute number of CD45^+^ immune cells was slightly reduced in ARF6^f/f^ tumors, no significant difference was observed in the fractions of CD4^+^ T cells, CD8^+^ T cells, B cells, macrophages, NK cells, or dendritic cells between ARF6^WT^ and ARF6^f/f^ tumors (Figure S2A-D). Given that IFN signaling and TNFα signaling were increased in ARF6^f/f^ tumors (Figure 2A), we hypothesized that CD8^+^ T cells in ARF6^f/f^ tumors could have enhanced antitumor activity. Indeed, ARF6^f/f^ tumors had significantly higher percentages of IFNγ^+^ and granzyme B (GzmB^+^) CD8^+^ T cells (Figures 3A-B), demonstrating enhanced CD8^+^ T cell effector function, which may explain why tumorigenesis and progression were limited without ARF6. There was no significant difference in CD8^+^ T cell effector function in spleens of *Arf6*^f/f^ and *Arf6*^WT^ mice (Figure 3A-B), indicating a localized effect within the TIME. There was also no significant difference in PD-1+ CD8+ T cells in ARF6^f/f^ tumors compared to ARF6^WT^ tumors (Figure 3B). Interestingly, ARF6^f/f^ tumors showed a significantly lower percentage of FoxP3+ Tregs and a lower Treg/CD8 ratio (Figure 3C-D), possibly reflecting alleviation of immune suppression.

**Figure 3:**
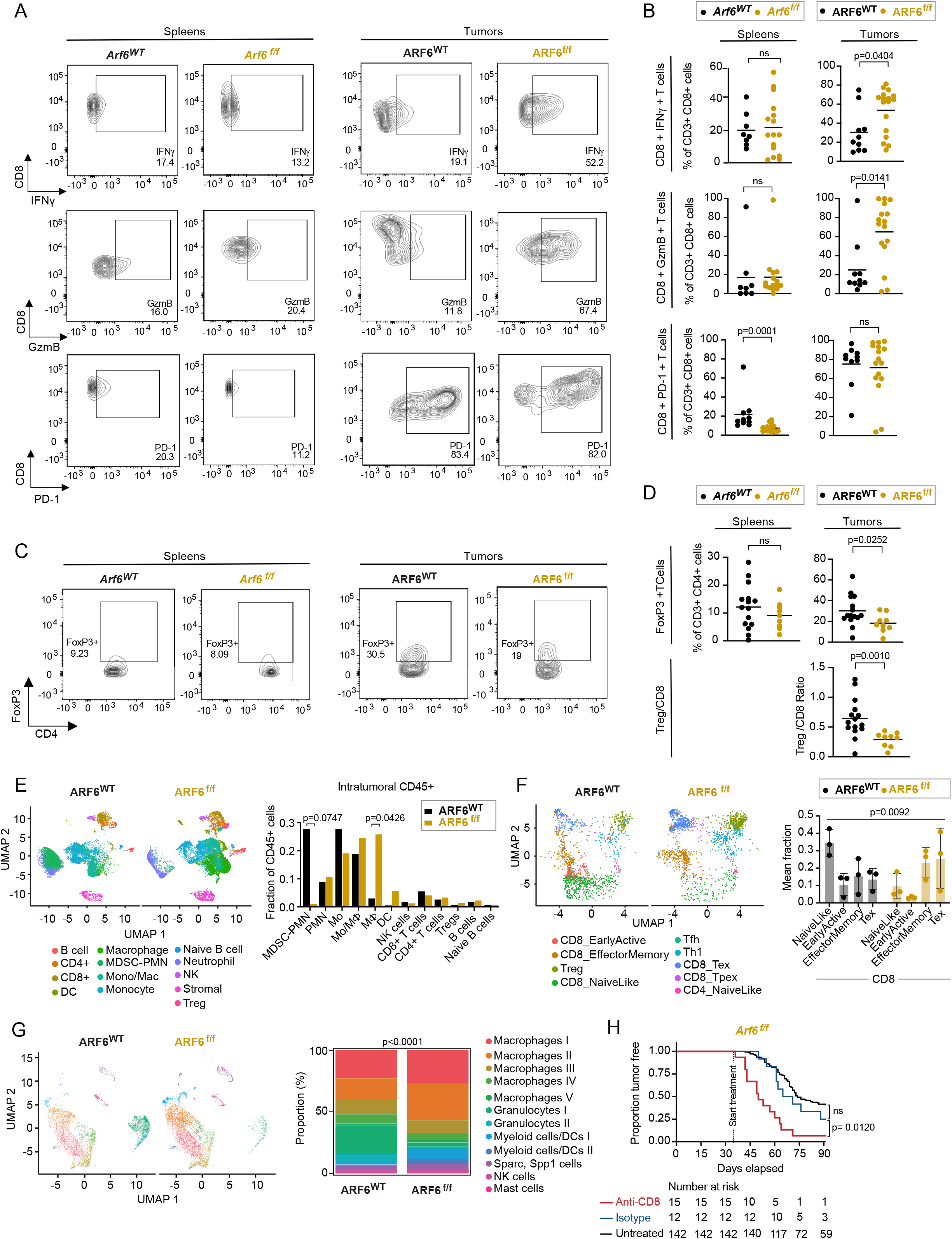
Heightened anti-tumor immunity in ARF6^f/f^ tumors. **(A and B)** Flow cytometry charts (A) and quantification (B) of IFNγ and granzyme B (GzmB) in CD8^+^ T cells from spleens and tumors of mice bearing ARF6^WT^ or ARF6^f/f^ tumors upon T cell reactivation. **(C and D)** Flow charts (C) and quantification (D) of the CD4+ FoxP3+ regulatory T cell fraction from spleens and tumors of mice bearing ARF6^WT^ or ARF6^f/f^ tumors. **(E)** Uniform manifold approximation and projection (UMAP) (scRNA-seq) showing intratumoral CD45+ cells (n= 19,367cells from ARF6^WT^, n= 28,003 cells from ARF6^f/f^ tumors, n=3 tumor each) and histogram showing mean % of immune cell types among total CD45+ cells. Unpaired t-tests. **(F)** UMAP showing projection of T cell clusters onto ProjecTIL reference and histogram showing mean % of CD8^+^ T cell subtypes among total CD8^+^ T cells. Likelihood ratio test, mixed effects model with fixed group effect and random effect for samples within groups. **(G)** UMAP showing tumor infiltrating myeloid cells and stacked histogram showing proportion of each cell type (%). Chi-square test of homogeneity. **(H)** Tumor-free survival of mice developing ARF6^f/f^ tumors without or with CD8^+^ T cell depletion. Antibody treatments were initiated when mice were 5-week-old and continued for 8 weeks. Kaplan-Meier log-rank test. See also Figure S2 and Figure S3.

To interrogate the TIME in greater detail, we subjected CD45+ tumor infiltrating immune cells to single-cell RNA sequencing (scRNA-seq) and found significant differences between genotypes (Figure 3E). ARF6^WT^ tumors contained a prominent population of polymorphonuclear neutrophil-derived, myeloid-derived suppressor cells (MDSC-PMN), defined by expression of *Cd84*, *Arg2*, *Irf1*, *Nfkbiz*, *Il1b*, *Csf1*, and *Ptgs2* ^35^. These MDSC-PMNs were notably absent from the ARF6^f/f^ tumors. In addition, there appeared to be a significant shift in the differentiation of monocytes into macrophages in the ARF6^f/f^ tumors. T cell clusters also showed significant differences between ARF6^WT^ tumors and ARF6^f/f^ tumors. Whereas naïve-like CD8^+^ T cells dominated ARF6^WT^ tumors, effector memory and cytolytic (exhausted) T cells dominated ARF6^f/f^ tumors (Figure 3F), concordant with an increased effector function of CD8^+^ T cells in ARF6^f/f^ tumors measured by flow cytometry (Figure 3A-B).

High proportions of CD11b^+^ myeloid cells were found in both ARF6^WT^ and ARF6^f/f^ tumors (Figures S2C and 3G). Further analysis of this population revealed 12 clusters: five were macrophages, two each were granulocytes and myeloid/dendritic cells, and one each was Sparc^+^ Spp1^+^ cells, NK cells and mast cells (Figure 3G). Compared to ARF6^WT^ tumors, ARF6^f/f^ tumors exhibited an increased fraction of macrophage clusters (macrophages I to V) and a corresponding decreased fraction of granulocyte clusters (granulocytes I to II). Within the five macrophages clusters, expression of IFNγ-inducible genes, such as MHC class II and Fc gamma receptor, were higher in ARF6^f/f^ tumors (Figure S2E), which was consistent with the higher IFNγ production observed in CD8+ T cells from ARF6^f/f^ tumors (Figure 2A). Overall, there were no significant differences in the expression of efferocytosis genes in macrophages (Figure S2E). These data suggest that there was heightened antigen presentation and opsonic phagocytosis of apoptotic cells by macrophages in the ARF6^f/f^ TIME.

Both the flow cytometry (Figure 3A-D) and scRNA-seq (Figure 3F) findings indicated a heightened antitumoral immune response mediated by CD8^+^ T cells. To confirm that the growth of ARF6^f/f^ tumors was restricted by the adaptive immune response, we treated *Arf6*^f/f^ mice with anti-CD8 antibody to deplete CD8^+^ T cells. Treatment of *Arf6*^f/f^ mice with anti-CD8 resulted in efficient removal of CD8^+^ T cells in spleen and tumor tissues (Figure S3) and accelerated tumor progression (Figure 3H). These results confirmed that CD8^+^ T cells restricted tumor progression in *Arf6*^f/f^ mice and that ARF6 was critical for tumor-cell intrinsic suppression of the adaptive immune response.

### ARF6 controls tumor intrinsic IFNγ signaling through trafficking of the IFNγR

With clear evidence of a heightened adaptive immune response and IFNγ signaling in the immune compartment of ARF6^f/f^ tumors, we speculated that ARF6 may regulate tumor intrinsic IFNγ signaling. In principle, loss of ARF6 could alter endocytic trafficking of the IFNγ receptor, affecting the acute, cytotoxic impact on tumor cells and/or the chronic immunosuppressive role of IFNγ. To investigate these possibilities, we interrogated IFNγ induced JAK-STAT signaling *in vitro*. Early passage murine melanoma cells from ARF6^f/f^ tumors showed a significantly reduced JAK1 and STAT1 phosphorylation after IFNγ stimulation (Figure S4A), suggesting that tumor intrinsic IFNγ signaling relies upon ARF6. Given the critical role of ARF6 in endocytic trafficking we hypothesized that ARF6 controls the surface expression of the IFNγ receptor. Indeed, cell surface and total levels of IFNγR1 were significantly reduced in ARF6^f/f^ murine melanoma cells (Figure 4A, 4B). However, the *Ifn*γ*r1* mRNA level was similar between ARF6^f/f^ and ARF6^WT^ tumors (Figure. 4C).

**Figure 4:**
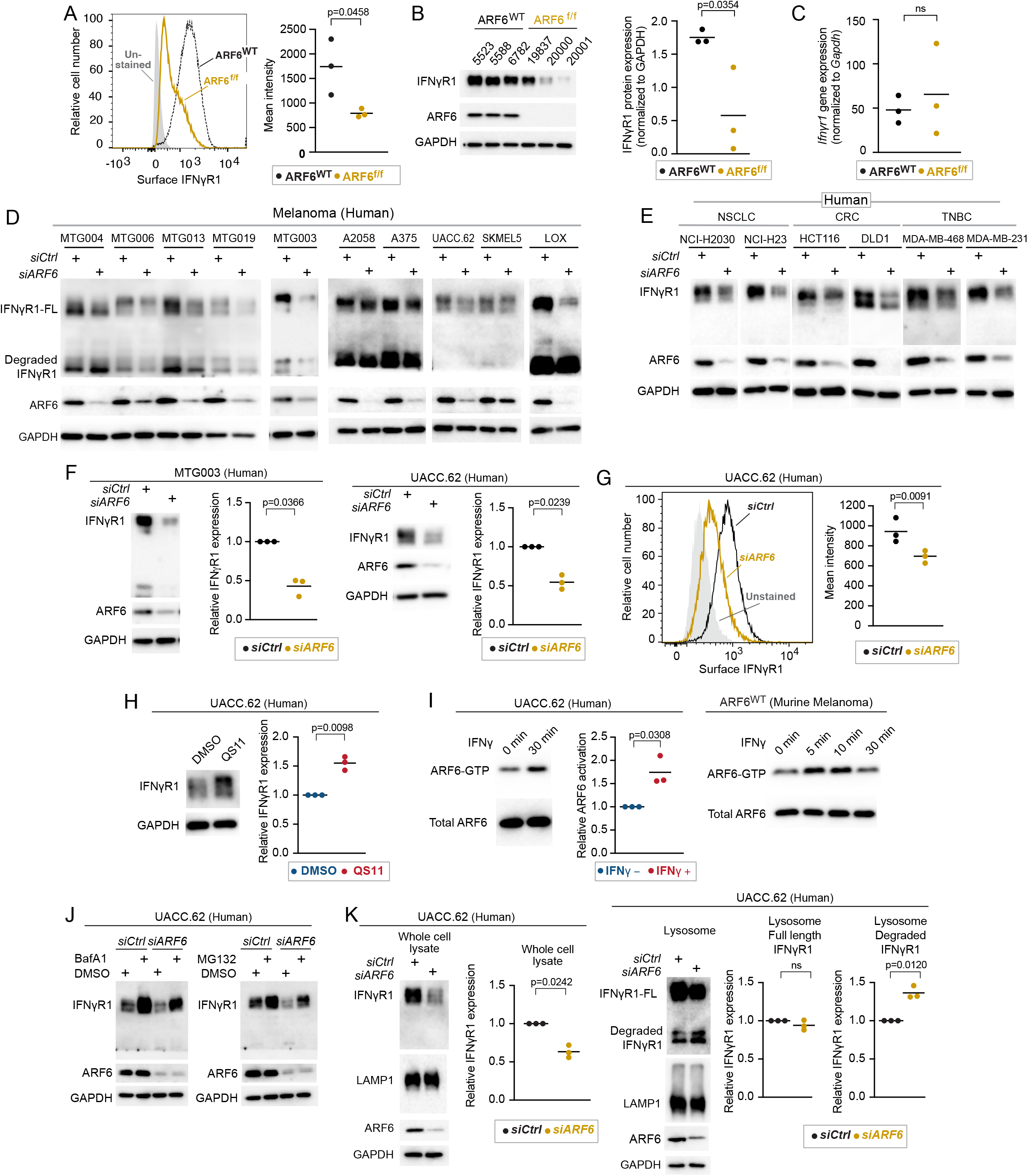
ARF6-dependent IFNγR1 surface expression in murine and human melanoma. **(A)** Flow cytometric detection of IFNγR1 cell surface expression in early-passage murine tumor cell lines, n=3 independent cell lines. **(B)** Western blot for IFNγR1 in early-passage murine tumor cell lines, n=3 independent cell lines. **(C)** Quantitative RT-PCR for IFNγR1 in one early-passage murine tumor cell line, n=3 replicates. **(D-F)** Western blot for full length (FL) and degraded IFNγR1, ARF6 and GAPDH in human melanoma patient derived melanoma cell lines (MTG) and commercially available melanoma lines (**D**), in human non-small cell lung cancer (NSCLC), colorectal cancer (CRC), triple negative breast cancer (TNBC) cell lines **(E)** Quantification of IFNγR1 in MTG003 (n=3) and UACC.62 (n=3) cells (**F**) without or with *ARF6* knockdown. **(G)** Flow cytometric detection of surface expression of IFNγR1 in UACC.62 cells without or with *ARF6* knockdown, n=3. **(H)** Western blot for IFNγR1 in UACC.62 cells without or with 2µM QS11 treatment for 24hours, n=3. **(I)** Western blot for total ARF6 and ARF6 GTP in UACC.62 cells without or with 500U/mL IFNγ treatment, n=3. **(J)** Western blot for indicated proteins in UACC.62 cells without or with ARF6 knockdown and without or with 50nM Bafilomycin A1 or 10µM MG132 treatment for 6h. **(K)** Western blot analyses of UACC.62 cells without or with ARF6 knockdown as indicated, n=3. **(A, B, C, G)** t-test. **(F, H, I, K)** Ratio paired t-test. See also Figure S4.

To test whether ARF6 controlled IFNγR1 protein levels in human tumors, we depleted ARF6 in early passage patient-derived melanoma cell lines and commercially available human melanoma cell lines. Knockdown of *ARF6* reduced the total IFNγR1 protein level in all of human melanoma cells tested (Figure 4D). Next, we asked whether this phenomenon was true in other cancers where ICB is a standard of care therapy, i.e., cancers that rely on IFNγ-driven AIR and where IFNγR density at the cell surface might impact therapeutic outcome. Knockdown of *ARF6* in cell lines derived from non-small cell lung cancer (NSCLC), mismatch-repair deficient colorectal cancer (CRC) and triple negative breast cancer (TNBC) similarly diminished the IFNγR1 protein level (Figure 4E), suggesting that ARF6-dependent regulation of IFNγR1 is conserved across cancer types. The total IFNγR1 protein level may, in fact, be tightly linked to the expression level of ARF6, as shown in Figure 4F where partial knockdown of *ARF6* reduced the IFNγR1 protein by half in human melanoma cells. Consistent with murine cells, IFNγR1 localization at the cell surface in human tumor cells was diminished by *ARF6* depletion (Figure 4G). In contrast, activation of ARF6 with the ARF6 GAP inhibitor QS11 ^19,36^ (Figure 4H, S4B) was sufficient to increase the total IFNγR1 protein level. Similarly, ectopic expression of ARF6^Q67L^ was sufficient to increase the total IFNγR1 protein level (Figure S4C), consistent with the effect of QS11 (Figure 4H, S4B) and confirming a specific role for ARF6-GTP in augmenting IFNγR1 protein level. Together these data provide evidence that the plasma membrane density and the total protein level of IFNγR1 in cancer cells depend on ARF6 expression and activity.

ARF6 is activated by growth factor receptors^18,37 38 16,39,40 41^, WNT-Frizzled^15,17^, Interleukins^19^, Toll like receptors^20 21^ and numerous G-Protein Coupled Receptors, (reviewed in ^14^). Nevertheless, activation of ARF6 by IFN receptors has not been reported. Importantly, IFNγ treatment significantly increased ARF6-GTP levels in murine and human melanoma cells (Figure 4I). These data implicate ARF6 in a feedback loop that enhances IFNγR1 protein level, possibly by ARF6-mediated recycling of the receptor to the plasma membrane. Internalized plasma membrane proteins that are not recycled can be trafficked to the lysosome^9^. Thus, we hypothesized that loss of ARF6 would result in IFNγR1 trafficking to the lysosome for degradation. To explore this possibility, we first examined how IFNγR1 was degraded in melanoma. Inhibition of either lysosomal or proteasomal degradation increased the total amount of IFNγR1 protein and partially restored the IFNγR1 level upon the depletion of ARF6 (Figure 4J). Hence, IFNγR1 protein stability is regulated by distinct mechanisms that may serve different cellular functions in tumor progression. Importantly, silencing *ARF6* led to significant enrichment of degraded IFNγR1 in the lysosomes (Figure 4K). Despite the well-known role of ARF6 in endocytosis, this increase in lysosomal degradation suggested that endocytosis of IFNγR1 was largely intact when ARF6 was depleted. Thus, our data suggest that ARF6 activation is essential for recycling, which is consistent with ARF6-mediated endocytic recycling of other receptors, such as the transferrin receptor ^42^ and the neurotrophic tyrosine kinase receptor type 1 (TRK1/TRKA)^43^. Overall, our data demonstrated that ARF6 prevented IFNγR1 from lysosomal degradation, suggesting that ARF6 may control how tumor cells respond to IFNγ in the TIME.

### IFNγ-driven Adaptive Immune Resistance Requires ARF6

Given that the net effect of tumor-specific deletion of *Arf6* was CD8^+^ T cell-mediated restriction of tumor progression, we interrogated the immunosuppressive output of IFNγ signaling in tumor cells. First, we used immunohistochemistry (IHC) to analyze the PD-L1 expression *in situ*. PD-L1 was present in a heterogeneous, multifocal pattern in the majority (70%) of ARF6^WT^ tumors tested whereas it was undetectable by IHC in ARF6^f/f^ tumors (Figure 5A), despite heightened IFNγ signaling detected in the TIME (Figure 2A). Both ARF6^WT^ and ARF6^f/f^ murine melanoma cells increased the expression of total and cell surface PD-L1 after exposure to IFNγ but ARF6^f/f^ cells expressed significantly less of both (Figure 5B-C). *CD274* (PD-L1) mRNA expression has been reported to peak between 6-12 hours after the start of IFNγ treatment in human lung cancer cells ^44^. In keeping with this time course, expression of *Cd274* was readily detectable in murine melanoma within 8 hours of IFNγ treatment (Figure 5D). Although ARF6 could potentially control the trafficking of PD-L1^23^, *Cd274* expression after IFNγ exposure was ARF6-dependent (Figure 5D). This can be explained by the diminished IFNγ-JAK-STAT signaling observed in ARF6^f/f^ melanoma cells (Figure S4A). In addition to PD-L1, ARF6^f/f^ tumor cells expressed lower levels of CD80, a CTLA-4 ligand, before and after IFNγ treatment, compared to ARF6^WT^ cells (Figure. 5F). Likewise, IFNγ-induced Indoleamine 2, 3-dioxygenases (IDO1) protein and mRNA expression were compromised by deletion or silencing of ARF6 (Figure 5F,G,H). IDO1-dependent catalysis of tryptophan generates kynurenine, inducing immunosuppressive Tregs^45^, as well as recruiting and activating MDSCs ^46^. Thus, reduced IFNγ-induced IDO1 could explain why there are significantly less Tregs and MDSCs in ARF6^f/f^ tumors (Figures 3D-E). In addition to immunosuppressive genes, IFNγ can also induce MHC Class I expression in melanoma to enhance tumor antigen presentation and immunogenicity ^47,48^. IFNγ treatment raised the level of MHC Class I protein, on the surface ARF6^f/f^ tumor cells, similar to that of unstimulated ARF6^WT^ tumor cells (Figure S5A), which appeared to be sufficient for antigen presentation and formation of MHC Class I – T cell receptor synapses based on the anti-tumor CD8 activity observed in the *Arf6*^f/f^ mice (Figures 3A, B, F, H). In contrast, expression of Gal3 and LSECtin, which are ligands of the immune checkpoint receptor LAG3 and are not IFNγ-inducible genes, were not dependent on ARF6 (Figures S5B-S5C). Thus, ARF6-dependent expression of immunosuppressive genes is not a generalizable phenomenon and may be limited to IFNγ-driven AIR.

**Figure 5:**
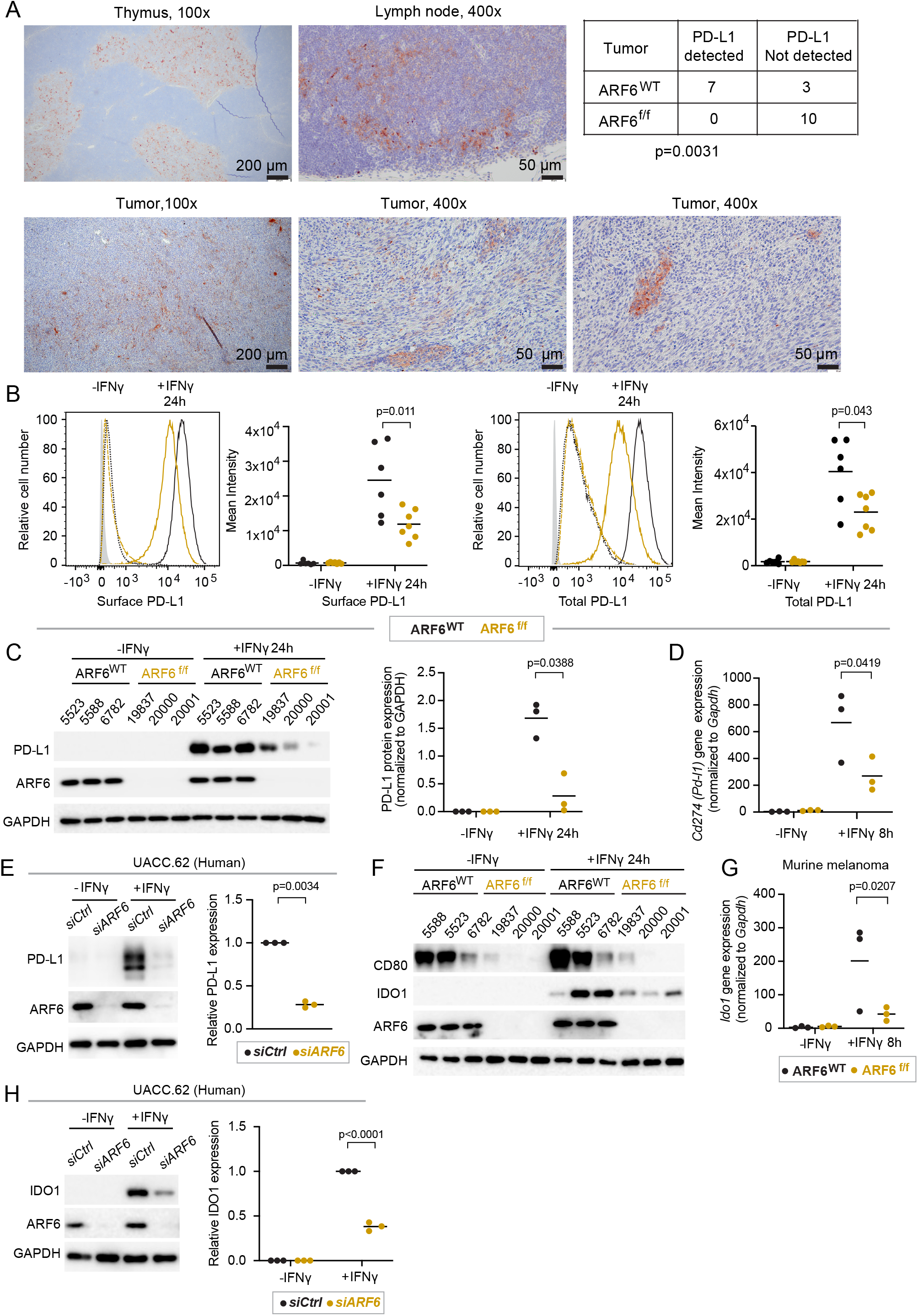
Activation of ARF6 and expression of immunosuppressive genes downstream of IFNγ. **(A)** Representative images of immunohistochemical (IHC) detection of PD-L1 in ARF6^WT^ tumors and summary of PD-L1 detection in n=10 tumors of each genotype tested, two-sided Fisher’s exact test. Thymus and lymph node are used as controls, **(B-D)** IFNγ-induced PD-L1 expression in early-passage murine tumor cell lines. **(B)** Flow cytometric detection of tumor cell surface and total protein, n=6-7 independent cell lines of each genotype. **(C)** Western blot for indicated proteins, n=3 independent cell lines of each genotype. **(D)** Quantitative RT-PCR analysis for *Cd274* mRNA, one tumor cell line of each genotype, n=3 replicates each. **(E)** Western blot for indicated proteins in UACC.62 cells, n=3. **(F)** Western blot for indicated proteins in early-passage murine tumor cell. **(G)** Quantitative RT-PCR for *Ido1* mRNA in one tumor cell line of each genotype, n=3 replicates each. **(H)** Western blot for indicated proteins in UACC.62, n=3 biological replicates. **(B, C, D, G)** Two-way ANOVA with Tukey’s multiple comparisons test. **(E, H)** Ratio paired t-test. See also Figure S5.

### Tumors with low ARF6 expression are insensitive to ICB

Given that IFNγ-induced PD-L1 mRNA and protein expression was dependent on ARF6, we hypothesized that anti-PD-1 treatment would fail to control tumor growth in *Arf6*^f/f^ mice. To test this, we treated *Arf6*^WT^ and *Arf6*^f/f^ mice with systemic anti-PD-1 antibody just prior to palpable tumor onset, when microscopic disease is expected. Anti-PD-1 treatment significantly limited tumor development in *Arf6*^WT^ mice (Figure 6A). Tumor growth was also restricted in *Arf6*^WT^ mice when treatment was initiated after tumors were well established (Figure S6A), confirming that in our model, tumor progression is partly reliant on PD-L1-mediated immune suppression and recapitulates IFNγ-driven AIR in human cancers. In contrast, anti-PD-1 treatment fails to alter tumor development in *Arf6*^f/f^ mice, (Figure 6A). Thus, in our model, ARF6 is necessary for IFNγ-driven AIR.

**Figure 6:**
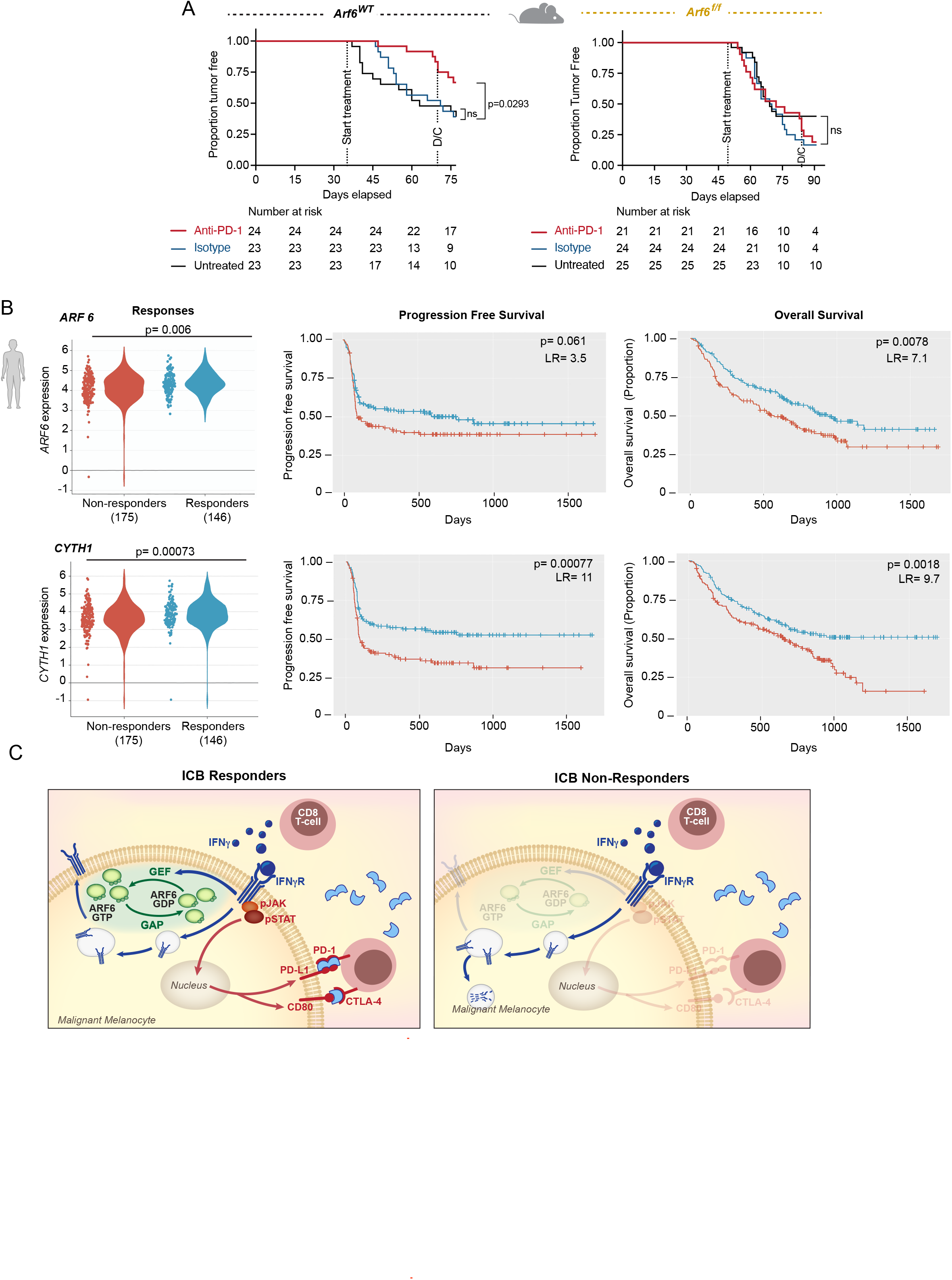
ARF6 status in tumors distinguishes ICB outcomes. **(A)** Systemic anti-PD-1 treated for 5 weeks duration. Includes mice that developed tumors within 35 to 77 (*Arf6*^WT^) days or 49 to 91 (*Arf6*^f/f^) days after Cre injection. Kaplan-Meier log-rank test. D/C= discontinued treatment. **(B)** Association of ICB treatment outcome with mRNA levels of *ARF6, CYTH1* in transcriptomes of pretreatment melanoma biopsies (CancerImmu expression analysis, aggregated data from n=10 queried melanoma clinical studies, adjusted p-values, Benjamini and Hochberg procedure, LR=likelihood ratio with df=1). **(C)** Proposed model of ARF6-mediated rheostatic control of ICB treatment outcomes. See also Figure S6.

Next, we asked whether the expression level of *ARF6* in patient tumors might correlate with responses to ICB. Cancer-Immu analysis of integrated data from ten melanoma cohorts^49^ showed that the overall expression of *ARF6* in pretreatment tumor biopsies was heterogenous among patients with advanced stage melanoma treated with ICB, and that the level of *ARF6* in these tumors significantly correlated with ICB outcomes (Figure 6B). Specifically, patients whose tumors expressed low levels of *ARF6* had inferior overall survival after ICB compared to those whose tumors expressed high levels of *ARF6* (Figure 6B). This was also true for the ARF6 GEF *CYTH1* (Figure 6B), which similar to ARF6, localizes to the plasma membrane ^50^. Table S1 lists all ARFs, GAPs and GEFs interrogated. *CYTH4* expression also correlated with outcome (Figure S6B), although this GEF is specific for ARF1 and ARF5 rather than ARF6 and its expression is reported to be limited to leukocytes, including T cells^51^. Thus, *CYTH4* expression may reflect tumor infiltrating immune cells in these melanoma samples (Figure S6B). Within the ARF family, ARF1 has been reported to cooperate with ARF6 in some scenarios ^52^. Nevertheless, ARF1 does not localize to the plasma membrane like ARF6 ^50^, and therefore would not be expected to have a role in IFNγR signaling. Thus, it is pertinent that *ARF1* expression in melanoma did not associate with ICB treatment outcomes (Figure S6C).

Surprisingly, low expression of *ACAP1* in tumors also associated with inferior survival with ICB therapy (Figure S6D). *ACAP1* localizes to recycling endosomes and while reduced expression may enhance ARF6 activation during endocytic trafficking, reduced *ACAP1* expression, in primary (Figure 1A) and metastatic tumors^24^, was prognostic (correlated with inferior survival of ICB treatment-naïve patient cohorts^29,53^). Therefore, the *ACAP1* correlates we observed in the ICB-treated cohorts may also reflect a prognostic status of this gene. In contrast, *ARF6* and *CYTH1* were not prognostic in untreated (TCGA^53^) patients (Figure S6E): rather their expressions were predictive of ICB treatment response, (Figure 6B). Thus, downregulation of *ARF6* and *CYTH1* expression may yield relatively low, tumor intrinsic ARF6-GTP, restricting the amount of IFNγR1 that can populate the plasma membrane and transmit signal that leads to effective anti-tumor immunity with ICB (Figure 4C). Similarly, low expression of the IFNγ-inducible immunosuppressive genes, *CD274* and *IDO1*, was also predictive of inferior survival of ICB-treated patients (Figure S6F-G).

## Discussion

Here, we report that the expression level of the endocytic trafficking protein ARF6 determines the capacity of melanoma to adapt to CD8^+^ T cell-mediated immunity (Figure 3) and can control response to ICB therapy (Figures 6, S6). Our data reveal a mechanism whereby the cytoplasmic pool of the IFNγR can be diverted away from the lysosome (Figure 4K) by ARF6 and maintained at the plasma membrane (Figure 4A,G) in sufficient quantities to support tumor intrinsic IFNγ signaling (Figure S4A) and downstream expression of immunosuppressive genes (Figure 5). This mechanism helps explain why loss of *Arf6* restricts primary tumor development and growth (Figure 1C-E), by unleashing CD8^+^ T Cell anti-tumor function (Figure 3H).

Somatic, loss of function mutations and copy number loss of genes in the IFNγ pathway occur in melanomas resistant to ICB ^54,55^, supporting that tumor intrinsic IFNγ signaling is essential for ICB treatment efficacy. Nevertheless, these somatic events have been reported as infrequent and, despite significant effort, the identification of more common mechanisms of resistance to ICB has been elusive^56^. *ARF6* and *CYTH1* expression in tumors are heterogenous among patients prior to therapy (Figure 6B) and our data suggests that their expression levels significantly influence response to ICB (Figure 6A, B, C). Inferior ICB outcomes associated with ARF6^LOW^ tumors could be the result of combined effects of 1) resistance to the acute, anti-proliferative and pro-apoptotic effects of IFNγ secreted by cytotoxic T cells and 2) the PD-1 checkpoint pathway being less functional due to insufficient IFNγ-dependent PD-L1 expression by tumors cells. Hence, our work reveals an alternative mechanism to help explain how tumor intrinsic IFNγ signaling could be diminished in ICB resistant tumors, through downregulation of ARF6.

The machineries that control endocytic transport of the IFNγR1 are poorly understood. Preliminary studies in HeLa cells have shown that while internalization of the IFNγR is mediated by clathrin-dependent endocytosis, unlike the IFNα receptor, ligand binding of the IFNγR results in clustering of the receptor in plasma membrane lipid microdomains and STAT1 activation that is independent of the clathrin system^57^. Thus, the initiation of IFNγ signaling may precede endocytosis. How the IFNγR1 receptor returns to the plasma membrane after endocytosis is unknown. Our data shed new light on this process and implicate ARF6 in recycling of the receptor. Importantly, we show evidence that ARF6-dependent regulation of the IFNγR1 protein is a conserved mechanism across high-incidence cancer types (Figure 4D, E), including NSCLC, colorectal cancer, TNBC and melanoma. In NSCLC and TNBC, the amount of PD-L1 expression by tumor cells, when evaluated with FDA-approved companion diagnostic testing, predicts response to pembrolizumab and determines patient eligibility for therapy^58–61^. Based on our data, it is plausible that ARF6 controls IFNγ-induced PD-L1 tumor expression in these cancers and, like melanoma, tumor expression of ARF6 may be related to ICB treatment outcomes.

In this study, the first clue that tumor intrinsic ARF6 controlled the TIME came from an unbiased comparison of ARF6^WT^ vs. ARF6^Q67L^ and ARF6^f/f^ primary tumors (Figure 2A). The decreased expression of inflammatory signatures in bulk transcriptomes from ARF6^Q67L^ primary tumors suggested immune suppression, which may help explain the aggressive metastatic behavior in this model^24^. Mechanistically, both pharmacologic and genetic activation of ARF6 were sufficient to increase IFNγR1 protein (Figure 4H and S4B-C). While not tested in this study, our data suggest that ARF6-GTP augments IFNγ-driven AIR. More work is needed to confirm this and investigate whether targeted activation of ARF6 (with pharmacologic agonists) might enhance ICB efficacy. For now, cumulative *in vivo* data from our immunocompetent models has demonstrated that ARF6 supports distinct, complementary tumor cell functions that can lead to both primary tumor and metastatic progression. While activated ARF6 elicits invasive behavior through multiple mechanisms^15,16,24,62,63^, here we have shown that ARF6 also promotes immune suppression. Thus, our immunocompetent models support that together, ARF6-dependent invasion and immune suppression during early tumor formation promote metastatic behavior (shown previously^24^).

The ARF6-dependent IFNγR1 trafficking mechanism presented here has potentially broad implications for inflammatory signaling pathways and future development of immuno-therapeutics. Despite its known roles in both endocytosis and recycling of cell surface cargo, our findings suggest that ARF6 may play a more significant role in recycling of IFNγR1, rather than internalization. If ARF6 were necessary for IFNγR1 internalization, knockdown or deletion of ARF6 should result in increased IFNγR1 expressed on the cell surface and a possibly a reduction in lysosomal degradation of the receptor. Our results showed the opposite, whereby loss of ARF6 led to lower IFNγR1 level on the cell surface (Figure 4A), and a concomitant increase of degraded IFNγR1 in the lysosome (Figure 4K). This is a compelling distinction that requires further investigation and may be pertinent to other inflammatory receptors. To the best of our knowledge, our data reveal a previously unknown mechanism of IFNγR trafficking. In benign cells, ARF6 is activated by and coordinates signaling and functional output of Interleukin-1β ^19^ and Toll-like receptors ^20 21^. Whether ARF6 is critical for trafficking of these, and other inflammatory receptors, remains to be elucidated in both benign and malignant pathologies. In bulk tumor transcriptomes, we detected ARF6-dependent IFNα and TNFα signaling together with IFNγ (Figure 2A). Although not tested in our study, it is possible that, like the IFNγR, ARF6 is activated by and traffics the IFNα and TNFα receptors to regulate their signaling. In the acute setting, ARF6 recycling of cytokine receptors might enhance CTL-mediated tumor killing. With chronic exposure, IFNγ and IFNα can cause epigenetic changes in cancer cells that lead to acquired resistance to ICB ^64^. Thus, more work is needed in understanding how ARF6-mediated endocytic trafficking may regulate inflammatory receptors, modulate the ability of cancer cells to adapt to immune attack and impact the efficacy of immunotherapy.

## MATERIALS AND METHODS

### KEY RESOURCES TABLE

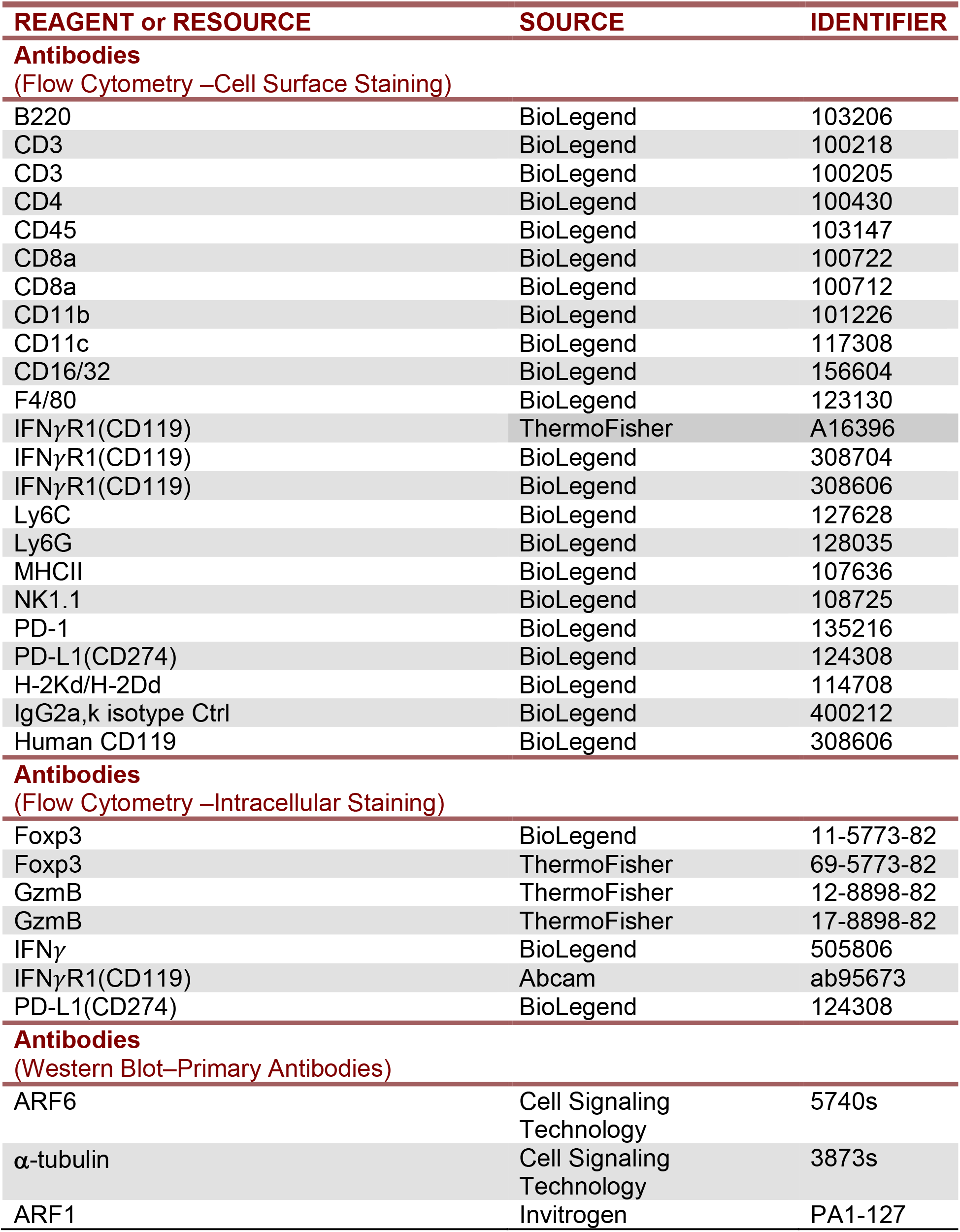
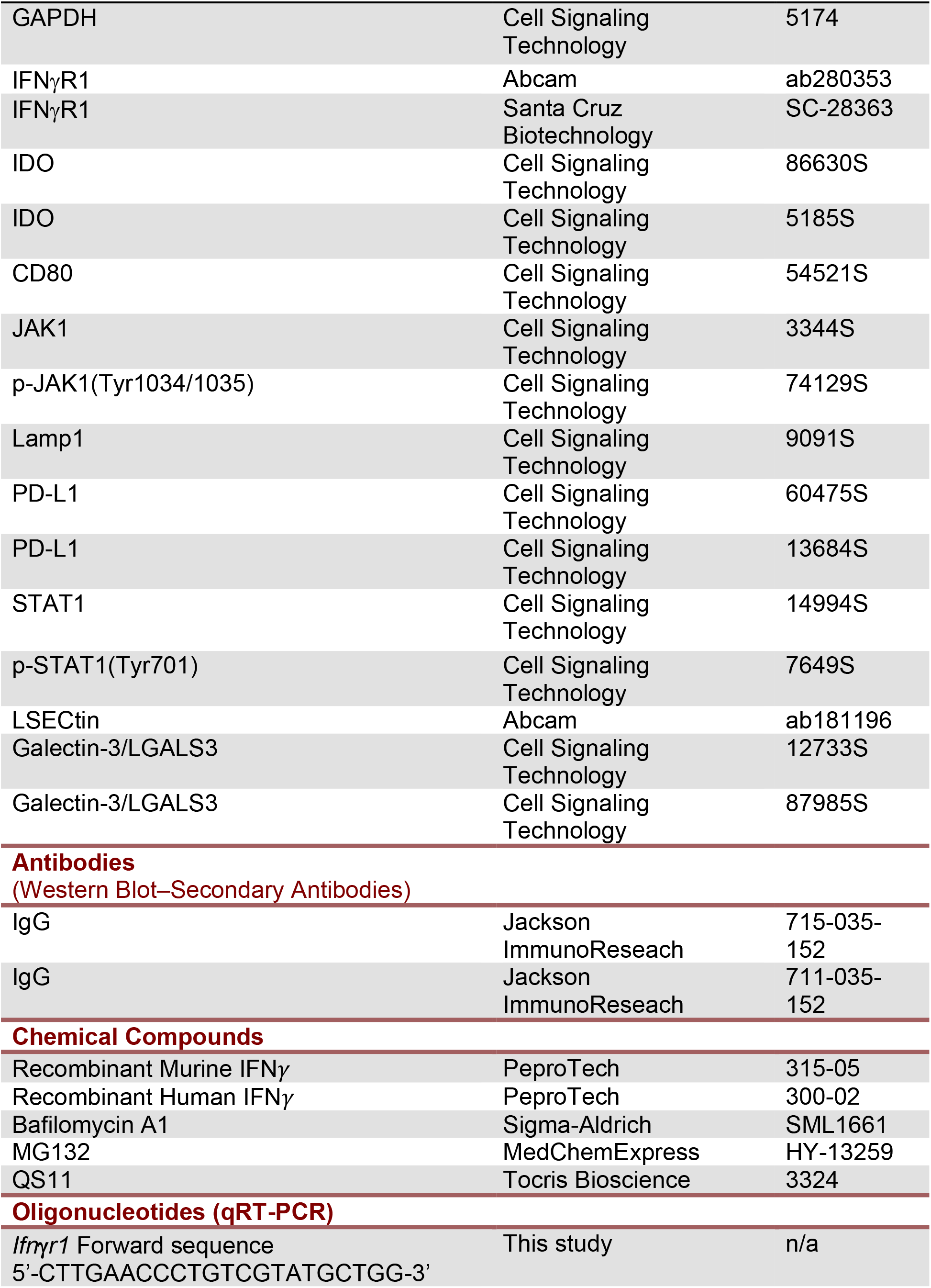
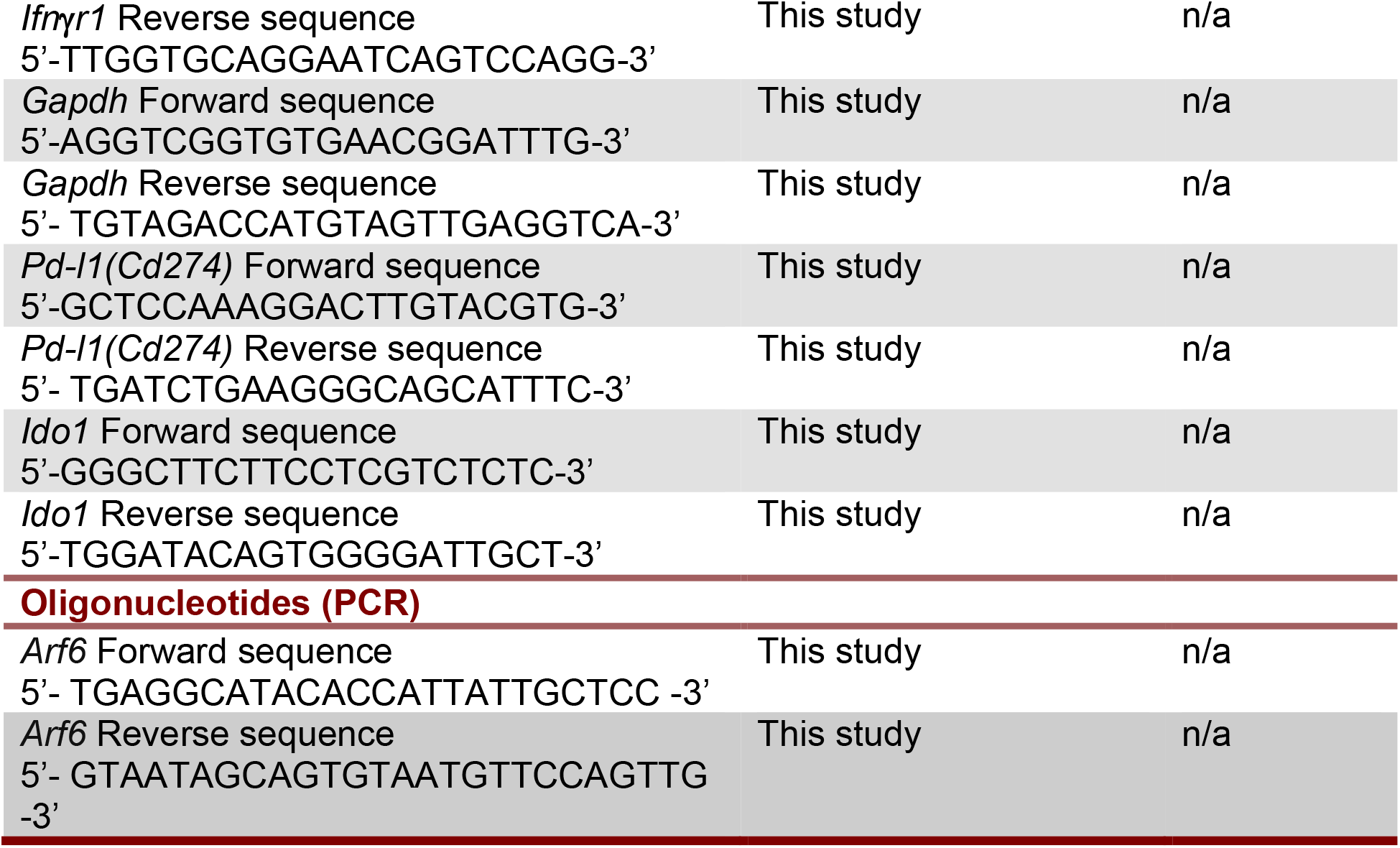

### EXPERIMENTAL MODELS

Animal studies were performed in accordance with a protocol approved by the University of Utah Institutional Animal Care and Use Committee. The *Dct::TVA*; *Braf^V^*^600^*^E^; Cdkn2a^f/f^* murine model was described previously^24,33^. Creation of the *Arf6^f/f^* allele was described previously^32^. The *Dct::TVA*; *Braf^V^*^600^*^E^; Cdkn2a^f/f^; Arf6^f/f^* mice were generated by backcrossing the *Arf6^f/f^* allele into *Dct::TVA*; *Braf^V^*^600^*^E^; Cdkn2a^f/f^* mice. Mouse colonies were maintained by random interbreeding. DF-1 cells infected with RCAS-Cre were suspended in HBSS (Gibco, Cat# 14025-092) and 50 µL of the cell suspension injected into the flank of neonate mice (0-3 days old) for two consecutive days. Tumor growth was measured by caliper every 1-3 days. Mice were euthanized once the primary tumor measured 2 cm in at least one dimension. Except where stated otherwise, non-tumor bearing mice were followed to 100 days before euthanasia. Tumor volume was calculated as the product of *length* x *width* x *depth* (mm^3^). Mice that did not develop tumors were excluded from onset and growth rate calculations.

Primary tumor cell lines from mice 5523, 5588, 6782, 7657, 21745, and 21793 were derived from primary melanomas harvested from *Arf6^WT^* mice. Primary tumor cell lines 19833, 19835, 19836, 19837, 19840, 19842, 19846, 19957, 19962, 20000, 20001, 20162, and 20163 were derived from *Arf6^f/f^* mice. Briefly, mechanically dissociated tumor cells were cultured with DMEM/F12 (ThermoFisher Scientific, Cat# 11330-032) supplemented with 10% v/v FBS (Atlas Biologicals, Cat# F-0500-DR), 0.5% v/v gentamicin (ThermoFisher Scientific, Cat# 15710072), 1% MEM Non-Essential Amino Acids Solution (ThermoFisher Scientific, Cat# 11140050) under standard conditions at 37°C in a humidified atmosphere with 5% CO_2_.

Human melanoma cell lines, A375, LOX-IMVI, SK-MEL-5, and UACC.62, were provided by Dr. M. VanBrocklin, Huntsman Cancer Institute (HCI). A2058 cells were purchased from the ATCC (Cat# CRL11147D). Early passage patient-derived human melanoma cell lines MTG003, MTG004, MTG006, MTG013, and MTG019, were provided by the Preclinical Research Resource (PRR) at HCI. A2058 and A375 cells were maintained in DMEM-high glucose (ThermoFisher Scientific, Cat# 11995073) supplemented with 10% v/v FBS, 1% v/v penicillin-streptomycin-glutamine. LOX-IMVI, SK-MEL-5, and UACC.62 cells were maintained in RPMI1640-high glucose media (ThermoFisher Scientific, Cat# A1049101) supplemented with 10% v/v FBS, 1% v/v penicillin-streptomycin-glutamine. Patient-derived human melanoma cell lines were maintained in Mel2 media provided by the PRR. Cells were incubated at 37°C in a humidified atmosphere with 5% CO_2_.

Non-small cell lung cancer cell lines, NCI-H2030, NCI-H23, were purchased from ATCC (Cat# CRL-5914 and CRL-5800) and maintained in RPMI1640-high glucose media supplemented with 10% v/v FBS, and 1% v/v penicillin-streptomycin-glutamine. Colorectal carcinoma cell line, HCT116, was purchased from ATCC (Cat# CCL-247). Colorectal carcinoma cell line, DLD-1, was provided by the PRR. HCT116 cells were maintained in McCoy’s 5A Modified media supplemented with 10% v/v FBS, and 1% v/v penicillin-streptomycin-glutamine. DLD-1 cells were maintained in RPMI1640-high glucose media supplemented with 10% v/v FBS, and 1% v/v penicillin-streptomycin-glutamine. Breast adenocarcinoma cell line, MDA-MB-468, was provided by the PRR. Breast adenocarcinoma cell line, MDA-MB-231, was purchased from ATCC (Cat# HTB-26). MDA-MB-468 and MDA-MB-231 cells were maintained in Leibovitz’s L-15 media supplemented with 10% v/v FBS, and 1% v/v penicillin-streptomycin-glutamine. Cells were incubated at 37°C in a humidified atmosphere with 5% CO_2_.

Human cell line authentication was performed at the University of Utah Genomics core facility using the Promega (Madison, WI) GenePrint 10 system.

DF-1 and A375-TVA cells were provided by Dr. S. Holmen (HCI). DF-1 cells were maintained in DMEM-high glucose supplemented with 10% FBS, 0.5% v/v gentamicin, and maintained at 39°C, with 5% CO_2_. A375-TVA cells were maintained in DMEM-high glucose supplemented with 10% FBS, 0.5% v/v gentamicin, at 37°C with 5% CO_2_ and were used to verify RCAS/Cre expression in DF-1 cells.

## METHOD DETAILS

### RNA interference

Silencing of endogenous *ARF6* in human cell lines was performed by sequential transfection of either *ARF6* siRNA (Qiagen, Cat# 1027417; GeneGlobe S02757286), and compared to AllStars Negative Control siRNA (Qiagen, Cat# 1027181) at a final concentration of 40 nM using Lipofectamine™ RNAiMAX transfection reagent (ThermoFisher Scientific, Cat# 13778150). Briefly, cells were seeded in a 6-well plate and first transfected with 40 nM siRNA mixed with 7.5 µL of Lipofectamine™ RNAiMAX transfection reagent. After 24 hours, transfections were repeated under the same conditions. Cells were collected 24 hours after the second transfection for flow cytometry and western blot analyses.

### Histology

Mouse tissues were fixed in 10% neutral buffered formalin overnight then placed in 70% ethyl alcohol before paraffin-embedding (FFPE). Four-micron sections of primary murine tumors were assessed by a board-certified pathologist (A.H.G.) blinded to the genetic identity of each sample for evaluation of PD-L1 immunohistochemical staining and for evaluation tumor-infiltrating lymphocyte clusters.

### Flow cytometry analysis

Tumors and spleens were harvested after euthanasia. Tumors were minced into small fragments and incubated with GentleMACS C tubes (Miltenyi Biotec, Cat# 130-093-237) with a serum free-DMEM/F12 solution containing digestive enzymes from a Mouse Tumor Dissociation Kit (Miltenyi Biotech, Cat# 130-096-730). Tumors were disaggregated into single-cell suspensions using a Miltenyi GentleMACS Octo Dissociator (Miltenyi Biotec). Tumor cells were filtered through a 70 μm nylon filter and treated with an RBC lysis solution (ThermoFisher Scientific, Cat# 00-4300-54). Spleens were disaggregated into single-cell suspensions by mechanical disruption and filtered through a 40 µm nylon filter and treated with an RBC lysis solution. 3-4 × 10^6^ cells from tumor or spleen were stained with antibodies against cell surface antigens for 1 hour on ice before flow cytometry analysis (BD Fortessa). For intracellular staining, cells were fixed and permeablized with a fixation/permeablization reagent (ThermoFisher Scientific, Cat# 00-5523-00) according to the manufacturer’s instructions. Cells were stained with antibodies against intracellular antigens for 30 minutes on ice before flow cytometry analysis.

To assess IFNγ and granzyme B (GzmB) production, tumors and spleens were disaggregated into single-cell suspensions using the method described above. 1 × 10^6^ cells were seeded and added to a cell activation cocktail (BioLegend, Cat# 423303) and incubated at 37°C in a 5% CO_2_ incubator for 6 hours. Stimulated cells were collected and stained with antibodies against cell surface antigens. After fixation and permeabilization, cells were stained with antibodies for intracellular IFNγ or GzmB for 30 minutes on ice before flow cytometry analysis. Antibodies used for flow cytometry analysis are listed in Key Resources Table.

### Quantitative reverse transcription polymerase chain reaction (qRT-PCR)

Total RNA from six independent primary murine tumor cell lines, 3 ARF6^WT^ (5523, 5588, and 6782) and 3 ARF6^f/f^ (19837, 20000, and 20001), were treated for 8 hours with 500 U/ml murine IFNγ then collected in RNAlater (ThermoFisher Scientific, Cat# AM7024). RNA was extracted using an RNeasy Plus kit (Qiagen, Cat# 74034) according to the manufacturer’s instructions. Extracted RNAs from each sample were converted into cDNA using SuperScript IV VILO (SSIV VILO) Master Mix (ThermoFisher Scientific, Cat# 11756050). QRT-PCR was performed in triplicate for each sample using PowerUp™ SYBR™ Green Master Mix (ThermoFisher Scientific, Cat# A25780) and run using a QuantStudio™ 6 Flex Real-Time PCR System (ThermoFisher Scientific) on 96-well plates. Primers used for qRT-PCR are shown in Key Resources Table. The specificity of the amplicons was assessed by melting curve analyses. Relative mRNA expression of each gene was calculated using the number of cycles needed to reach the specific threshold of detection (CT) and normalized to the expression of *Gapdh*.

### Western blot and ARF6 GTP-pulldown

Cells were lysed using Pierce® IP Lysis buffer (ThermoFisher Scientific, Cat # 87788) with 1X Halt™ Protease and Phosphatase Inhibitor Cocktail (ThermoFisher Scientific, Cat# 78442). Protein concentrations were determined using the Pierce™ BCA Protein Assay Kit (ThermoFisher Scientific, Cat# 23227). Cell lysates were boiled with SDS sample buffer. Proteins from the cell lysates were separated by SDS polyacrylamide gel electrophoresis (SDS–PAGE) and transferred to polyvinylidene difluoride (PVDF) membranes (ThermoFisher Scientific, Cat# 88518). The PVDF membranes were blocked with TBST (10 mM Tris-HCl, 150 mM NaCl and 0.1% v/v Tween-20) containing 5% v/v skim milk and incubated with primary antibodies. After washing in TBST, membranes were incubated with HRP-conjugated secondary antibodies then washed with TBST before developing with Western Lightning™ Plus Chemiluminescence Reagent (PerkinElmer, Cat# NEL103001EA) or SuperSignal™ West Dura Extended Duration Substrate (ThermoFisher Scientific, Cat# 37075). Luminescent signal was detected using the Azure c300 or c600 (Azure Biosystems). ImageJ (NIH, Bethesda, MD, USA) was used to quantify the intensity of bands on the blots. Images were adjusted equally for brightness and contrast using ImageJ or Adobe Photoshop (Adobe Inc.). Antibodies used for western blots are listed in Key Resources Table.

ARF6-GTP pull-downs were performed using GGA3 PBD Agarose beads (Cell Biolabs, STA-419) as previously described^22^. Briefly, cells were starved in 0.1% FBS for 16h before treatment with IFNγ and/or QS11. After treatment, cells were lysed with ARF6-pulldown lysis buffer (Cell Biolabs, Cat# 240102) including 1X Halt™ Protease and Phosphatase Inhibitor Cocktail. Lysates were centrifuged, and supernatants were added to GGA3-conjugated beads and agitated for 1 hour at 4°C. Beads were washed in ARF6-pulldown lysis buffer and prepared for western blot analysis.

### *Arf6* mRNA *in situ* hybridization

*Arf6* mRNA was detected in four μm tissue sections using a custom *Arf6* probe (Cat# 1205481-C1, Advanced Cell Diagnostics), targeting a sequence located between the loxP sites of the *Arf6^f/f^* allele, and the RNAscope 2.5 HD Reagent Kit - RED (Cat# 3222350, Advanced Cell Diagnostics) according to the manufacturer’s instructions.

### Lysosome enrichment

RNA interference was performed on UACC.62 cells as described above. Lysosome enrichment was performed using a Lysosome Enrichment Kit (ThermoFisher Scientific, Cat# 89839) according to the manufacturer’s instructions. Briefly, cells were disrupted in the supplied lysis buffers using a Dounce homogenizer. Following centrifugation to pellet debris, supernatants were loaded on 15%-30% Optiprep (Millipore-Sigma, Cat# D1556) step gradients and centrifuged at 145,000 g for 2 hours. The lysosome-enriched fractions were collected and pelleted prior to lysis and quantitation for use in western blot analysis.

### *In vivo* CD8+ T-cell depletion

*Arf6^f/f^* mice at 5 weeks post DF1 RCAS-Cre injection were treated with antibody prior to tumor onset. Anti-CD8 (200 µg/mouse, Bio X Cell, Cat# BE0117) or rat IgG2b isotype control (200µg/mouse, Bio X Cell, Cat# BE0090) were injected intraperitoneally twice per week for 8 weeks or until the tumor measured 2 cm in one dimension. Mice were euthanized once the primary tumor measured 2 cm in at least one dimension, or 1 week after the conclusion of treatment. Tumor incidence and onset day were documented. CD8+ cell depletion was verified by flow cytometry.

### *In vivo* anti-PD-1 treatment

Anti-PD-1 treatment was initiated prior to tumor onset in *Arf6^WT^* mice at 5 weeks, and in *Arf6^f/f^* mice at 7 weeks. Anti-PD-1 (8 mg/kg, Bio X Cell, Cat# BE0146) was administered intraperitoneally twice per week for 5 weeks. A second cohort of *Arf6^WT^* and *Arf6^f/f^* mice containing established tumors was treated with Anti-PD-1 when tumors measured approximately 0.4 – 0.5cm in greatest dimension. Anti-PD-1 was administered at 8 mg/kg and mice were euthanized once the primary tumor measured 2 cm in one dimension.

### Proteomics

Protein extraction and reverse-phase protein array of frozen mouse tumors was performed by the MD Anderson Cancer Center Functional Proteomic RPPA Core Facility.

### Single-cell RNA sequencing and analysis

Tumors were dissociated as described above under flow cytometry analysis. Resulting cells from tumors were stained with antibodies against CD45 and sorted (FACSAria 5 Laser). CD45+ cells were then subjected to single-cell RNA sequencing (scRNAseq) analysis using the 10X Genomics Chromium system and run on an Illumina NovaSeq instrument by the High Throughput Genomics core (University of Utah).

The Fastq files were aligned to the mm10 mouse reference from 10X genomics (refdata-gex-mm10-2020-A) using cellranger count (version 6.0.1) to create quality control metrics, Loupe Browser files, and filtered gene barcode matrices^65^. The filtered gene barcode matrices were loaded into the Seurat 4.1.1 package and merged into a single matrix^66^. About 5% of cells with more than 20% mitochondrial reads, fewer than 200 features, or more than 8000 features were removed and counts were normalized using the sctransform method^67^. Twenty dimensions and 0.8 resolution were selected for clustering and the non-linear dimensional reduction with UMAP. Cluster markers and differentially expressed genes were identified using the Wilcoxon Rank Sum test in Seurat. The list of cluster markers and significant genes were analyzed using the Enrichr website to identify over-represented gene sets, pathways and cell types using a Fisher’s Exact Test^68^. Cell type labels were assigned to cells using a nearest neighbor classifier in the SingleR package^69^. T cell clusters were then projected onto a T cell reference using the ProjecTILs package^70^.

Macrophage single-cell RNA sequencing analysis was performed using the R package Seurat (version 4)^71^. Low quality cells (greater than 10% mitochondrial genes, fewer than 1000 or more than 5000 RNA features per cell) and lymphocytes (*Ighm* > 0.001 and *Cd3e* > 0.001) were filtered out. Gene expression values were normalized, scaled, and aligned across conditions. Then, cells were clustered and UMAP was used to visualize the distribution of these clusters. A cluster of cells enriched with *Ly6c1*, *Gpihbp1*, *Cd36*, and *podxl* genes, with extremely low frequencies (0.15% and 0.04% in ARF6^WT^ and ARF6^NULL^ tumors, respectively) of *Cd3e/Ighm* negative cells, were excluded from the UMAP. Differentially Expressed Genes (DEGs) for selected clusters were obtained using the Seurat function FindMarkers with the default parameters (non-parametric Wilcoxon rank sum test). To facilitate the exploration of genes of interest, the default logFC threshold was reduced, enabling the discovery of several smaller, yet significant, changes in gene expression that had been previously excluded.

### Bulk RNA sequencing and analysis

RNA was extracted from fresh-frozen mouse tumors using Direct-zol RNA Miniprep Plus Kit (Zymogenetics) after frozen section histologic confirmation of high tumor content. RNA-sequencing was performed on 6 tumors from each genotype (*Arf6*^WT^, *Arf6*^f/f^, and *Arf6*^Q67L^) using Illumina TruSeq Stranded mRNA Library Preparation Kit with polyA selection followed by lllumina HiSeq 2500 125-cycle paired end sequencing.

Bulk tumor RNA sequencing analysis included GSEA using MSigDB Hallmark (NES scores). The mouse GRCm38 genome and gene feature files were downloaded from Ensembl (release 90) and a reference database was created using STAR (version 2.5.2b) with splice junctions optimized for 125 base pair reads^72^. Reads were trimmed of adapters using cutadapt (version 1.16)^73^ and aligned to the reference database using STAR in two pass mode to output a BAM file sorted by coordinates. Mapped reads were assigned to annotated genes using featureCounts (version 1.6.3)^74^. The output files from cutadapt, FastQC, FastQ Screen, Picard CollectRnaSeqMetrics, STAR, and featureCounts were summarized using MultiQC to check for any sample outliers^75^. Differentially expressed genes were identified using a 10% false discovery rate with DESeq2 (version 1.26.0)^76^. Significantly enriched Hallmark, KEGG, and REACTOME pathways from MSigDB were detected using the fast gene set enrichment package^77^.

### Analysis of Leeds Melanoma Cohort

The normalized Leeds Melanoma Cohort gene expression dataset (EGAD00010001561) was downloaded from the European Genome-phenome Archive with permission from the University of Leeds, United Kingdom. The first seven columns in the "LMCFFPEmelanomanormalised.txt" file were used to create a survival table with stage, status, and overall survival time. The remaining columns included Illumina HT12.4 probes that were mapped to gene names using the illuminaHumanv4.db BioConductor package^78^. The log2 normalized counts from 40 ARF6-related genes were analyzed using a proportional hazards regression model using the coxph function in the survival package in R (version 3.5-5)^79^. In addition, gene expression within each AJCC stage class 1, 2, and 3 was modeled separately. Survival curves were plotted using high and low expression groups divided into both medians and quartiles.

### Cancer-Immu analysis

A pan-cancer analysis was performed on pre-treatment biopsy datasets using all samples from all ten melanoma study cohorts included in the Cancer-Immu Immunogenomic Atlas for Immune Checkpoint Blockade Immunotherapy^49^. Individual genes from the ARF6 pathway (Table S1), were queried with default parameters (median gene expression, sum cutoff of 0.5).

### TCGA analysis

The Cancer Genome Atlas (TCGA) melanoma RNA-Seq data were extracted from all melanoma TCGA RNA-Seq data (GDC Data Release 34.0 queried on July 27, 2022 in the TCGA_SKCM_v34.html). Survival times were generated using days to death and days to last follow up data. Samples without survival data were excluded. Each gene was evaluated individually and those with statistical significance. Samples were stratified into *ARF6* or *CYTH1* high vs. low groups by median centering of expression levels for each gene. Survival p values were calculated by log rank (Mantel– Cox) test.

### Quantification and Statistical Analysis

Statistical tests were assessed using Prism 8 or 9 software (GraphPad) or SAS version 9.4 (SAS Institute Inc). Quantitative values are represented as the mean of at least three replicates.

## Acknowledgements

We thank Diana Lim and Nikita Abraham for preparation of scientific graphics and illustrations; J.P. Snook for technical support; the Cell Response and Regulation (CRR) Program from the Huntsman Cancer Institute (HCI); HCI Shared Resources: Research Histology, Research Immunohistochemistry, High Throughput Genomics and Cancer Bioinformatics, Cancer Biostatistics and Preclinical Research Resource; University of Utah Flow Cytometry Core and the Genomics Core; MD Anderson Cancer Center Functional Proteomics Core, the Leeds Institute of Cancer and Pathology, University of Leeds, U.K.; Qi Liu, Hua-Chang Chen and Jing Yang (Vanderbilt University Medical Center) for Cancer-Immu technical support. This project was supported by funding from NIH/NCI P30CA042014 (HCI). A.H.G. has been supported by the American Cancer Society 133649-RSG-19-019-01-CSM and NIH/NCI K08CA188563, and is currently supported by the Department of Pathology at the University of Utah, NIH/NCI R37CA230630 and U.S. Department of Defense (DoD) W81XWH2210910. S.L.H. is supported by NIH/NCI R01CA121118. M.A.W. is supported by DoD W81XWH2210776. K.C.F. is supported by NIH R01AI158710. M.A.D. is supported by the Dr. Miriam and Sheldon G. Adelson Medical Research Foundation, the AIM at Melanoma Foundation, the NIH/NCI P50CA221703, the American Cancer Society and the Melanoma Research Alliance, Cancer Fighters of Houston, the Anne and John Mendelsohn Chair for Cancer Research, and philanthropic contributions to the Melanoma Moon Shots Program of MD Anderson.

## Author Contributions

Conceptualization, A.H.G, Y.W.; investigation, Y.W., J.W., E.C.W, C.P.R., A.R., E.D., J.K.H.T., R.K.W., A.H.G.; formal analysis, C.S., K.B. J.M.O, K.C.F., A.E.; methodology, S.L.H., K.C.F, M.A.W., A.H.G.; resources, Z.T., M.A.W., S.L.H., V.G.Y., M.A.D.; supervision, A.H.G., R.K.W., K.C.F., M.A.W., S.L.H.; writing-original draft, A.H.G, Y.W.; writing-reviewing & editing, A.H.G, Y.W., J.W., E.C.W., J.K.H.T., R.K.W.; funding acquisition, A.H.G.

---

**Figure S1:**
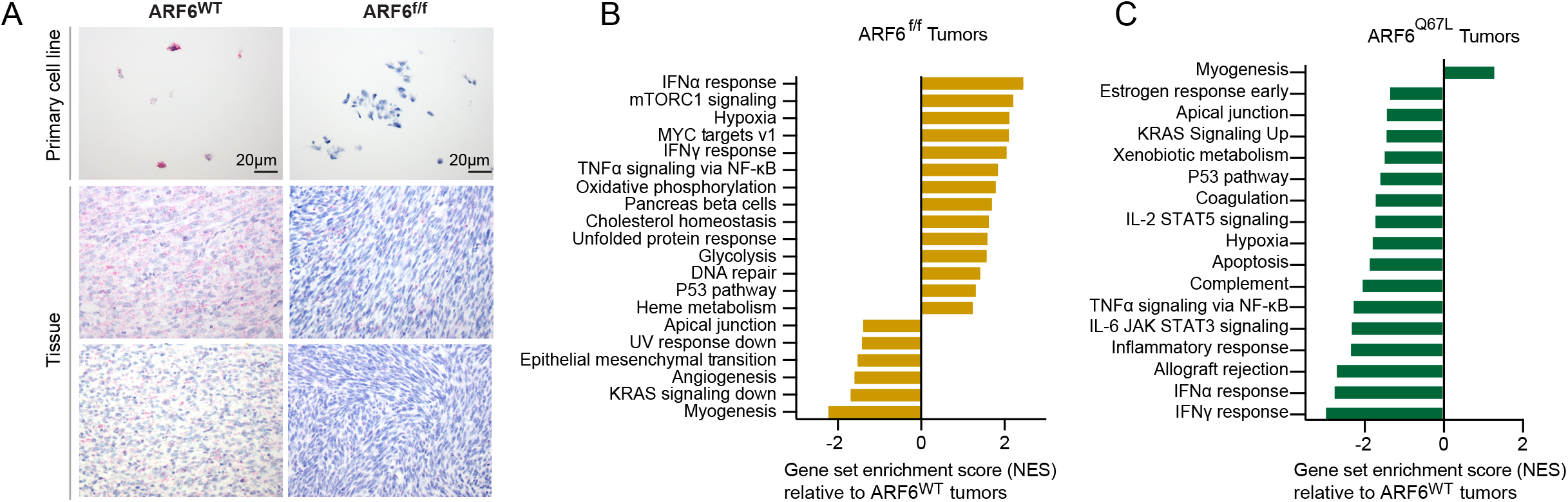
*Arf6* expression and ARF6-dependent gene expression pathways in murine tumors, related to Figures 1, 2. **(A)** In situ hybridization detection of *Arf6* mRNA (pink). Left panels show expected diffuse signal. Right panels show expected loss of signal. Right middle panel shows representative low-level heterogenous *Arf6* signal in murine tumor 19835, consistent with the low level of ARF6 detected by Western blot for the 19835 primary tumor cell line (see Figure 1G). **(B and C)** Bulk tumor transcriptomes (RNAseq) with significantly enriched gene sets (MSigDB Hallmark) in ARF6^f/f^ **(B)** and ARF6^Q67L^ **(C)** tumors compared to ARF6^WT^control tumors.

**Figure S2:**
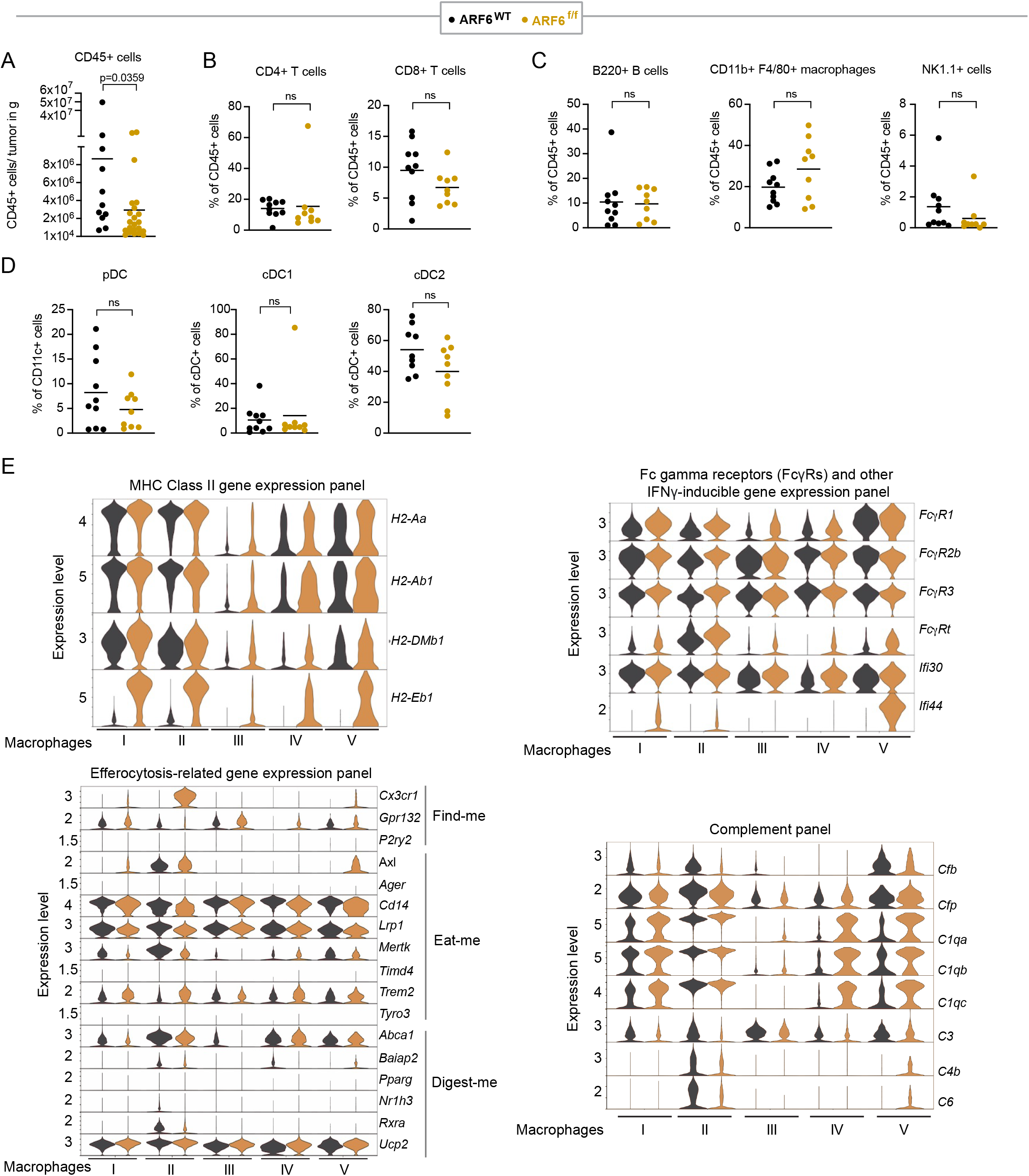
Immune profiling of tumor microenvironment, related to Figure 3. **(A)** The absolute numbers of CD45^+^cells per gram of tumor. **(B)** Fractions of CD4^+^and CD8^+^ T cells in CD45^+^cells. **(C)** Fractions of B220^+^B cells, CD11b^+^ F4/80^+^ macrophages, and NK1.1^+^ cells in CD45 ^+^cells. **(D)** Fractions of plasmacytoid dendritic cells (pDC) and conventional dendritic cell subsets (cDC1 and cDC2). **(A-D)** Graphs represent mean. Two-tailed Mann-Whitney t-test. **(E)** Expression of IFNγ-inducible genes related to antigen presentation (MHC Class II), phagocy-tosis (FcγR and other genes), efferocytosis-related genes and Complement genes, across different subtypes of macrophages. A comprehensive list of adjusted p-values, obtained from Seurat’s Wilcoxon Rank Sum test for differentially expressed genes, is provided in Table S2.

**Figure S3:**
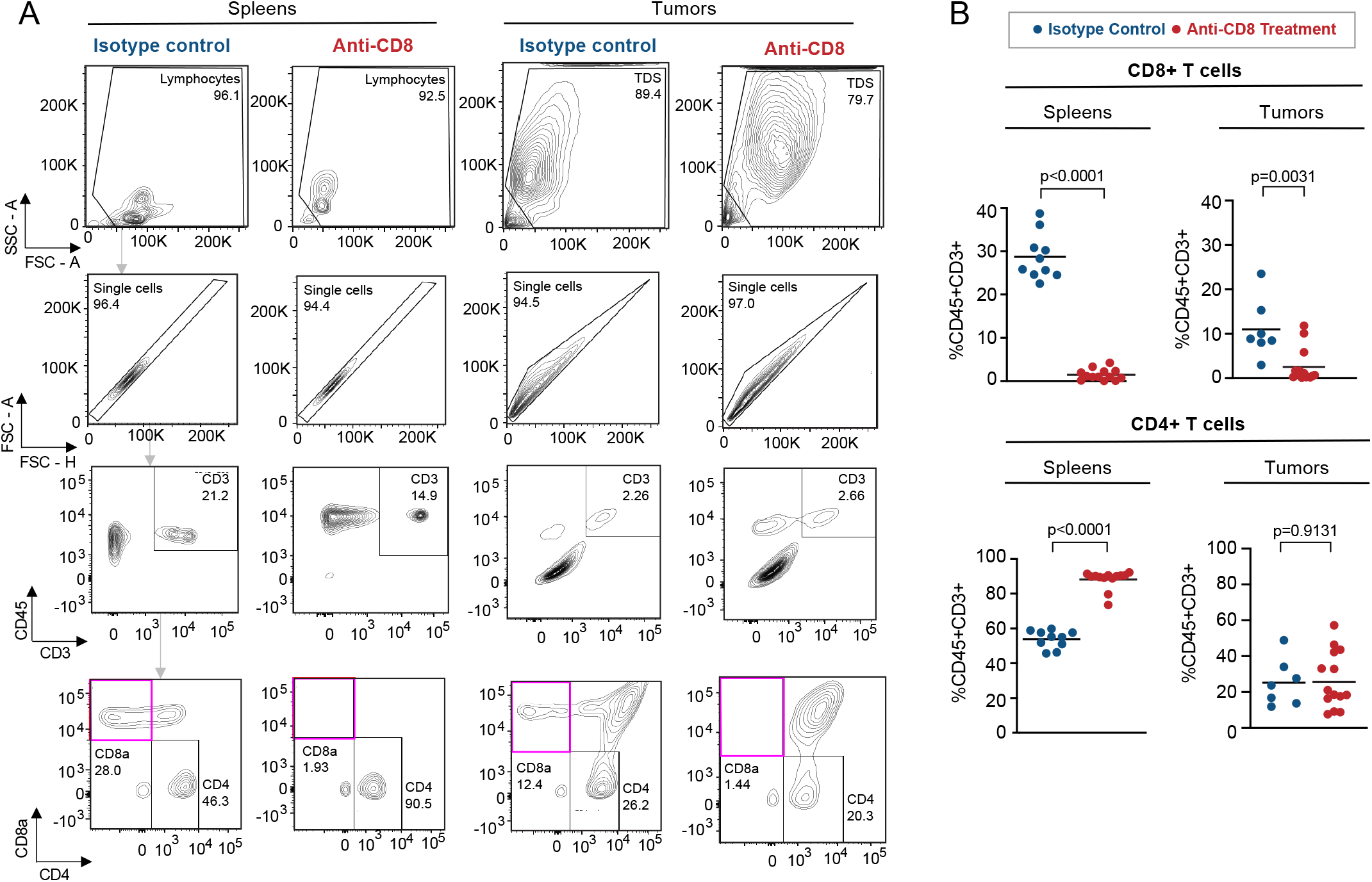
Efficiency of CD8 T cell depletion, related to Figure 3. **(A and B)** Quantitation of T cells by flow cytometry **(A)** and graph representing the mean **(B)** in spleens and tumors of mice treated with isotype control (IgG2b) or anti-CD8 antibody. Two-tailed Mann-Whitney t-test.

**Figure S4:**
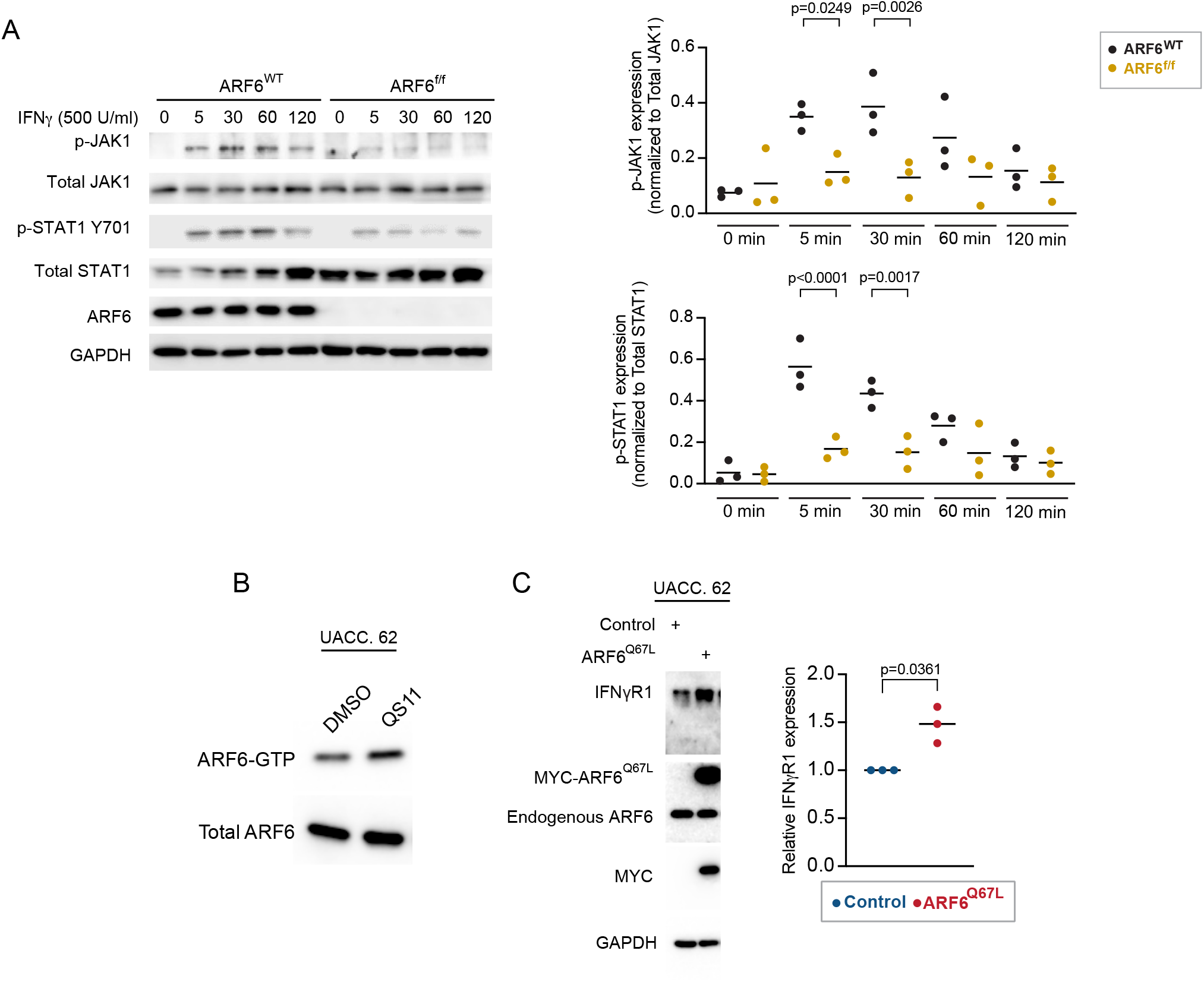
Tumor-intrinsic ARF6-dependent IFNγ signaling, related to Figure 4. **(A)** IFNγ-induced JAK-STAT signaling detection in early-passage murine melanoma cell lines. n=3 replicates each. Two-way ANOVA with Tukey’s multiple compari-sons test. **(B)** Total ARF6 and ARF6 GTP pulldown in UACC.62 cells without or with 2µM QS11 treatment for 1hr. **(C)** Western blot for indicated proteins in UACC.62 cells without or with adenoviral-mediated ectopic expression of constitutively active ARF6 (ARF6^Q67L^), control= empty vector, n=3 biological replicates. Ratio paired t-test.

**Figure S5:**
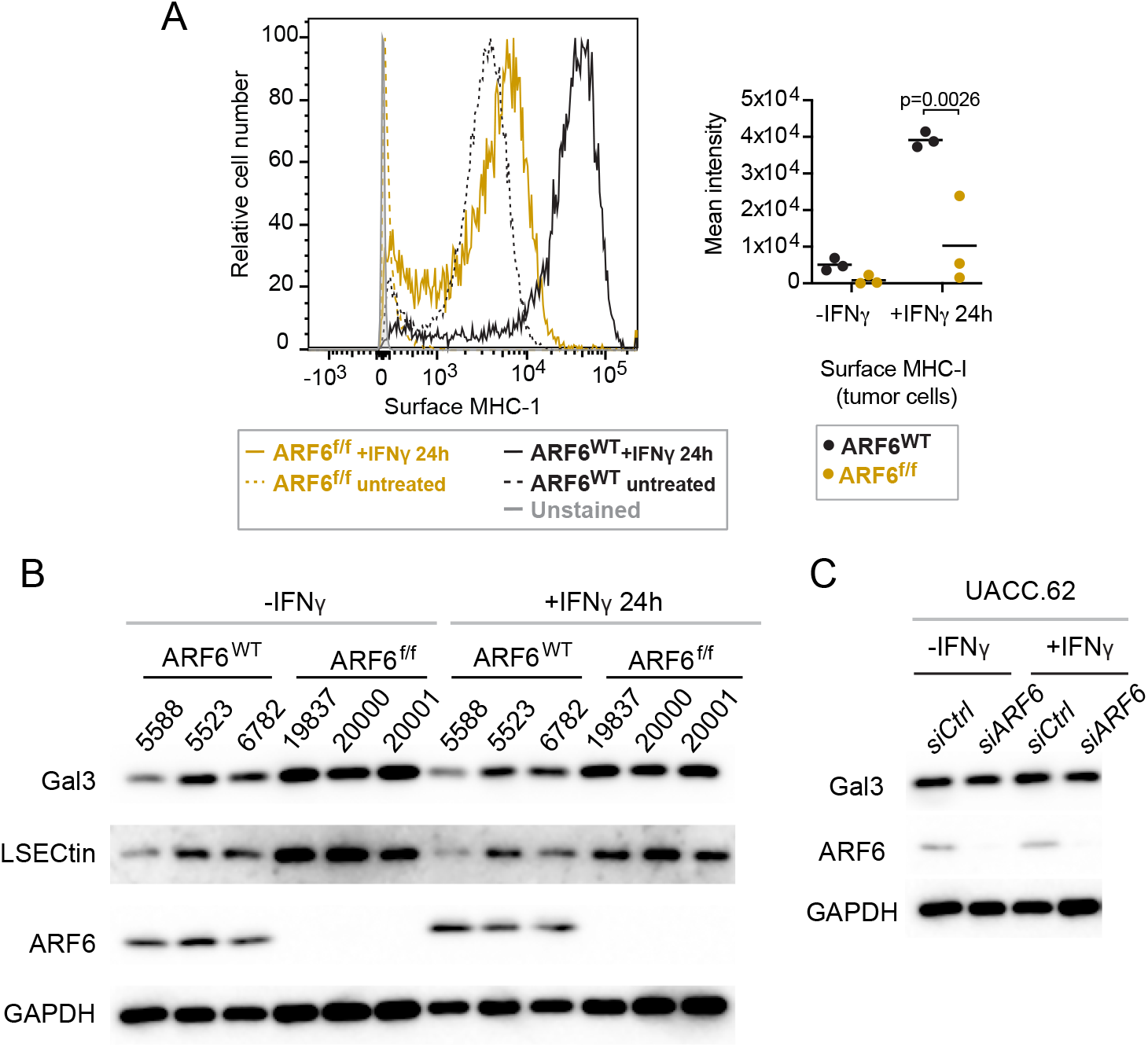
Expression of MHC-1 and LAG3 ligands, related to Figure 5. **(A)** Flow cytometric detection of tumor cell surface MHC-I expression, n=3 independent cell lines of each genotype. Two-way ANOVA test. **(B)** Western blot detection of Galectin3 (Gal3) and LSECtin in murine melanoma, n=3 independent cell lines of each genotype. **(C)** Western blot detection of Gal3 in UACC.62 cells without or with ARF6 knockdown, n=3 biological replicates.

**Figure S6:**
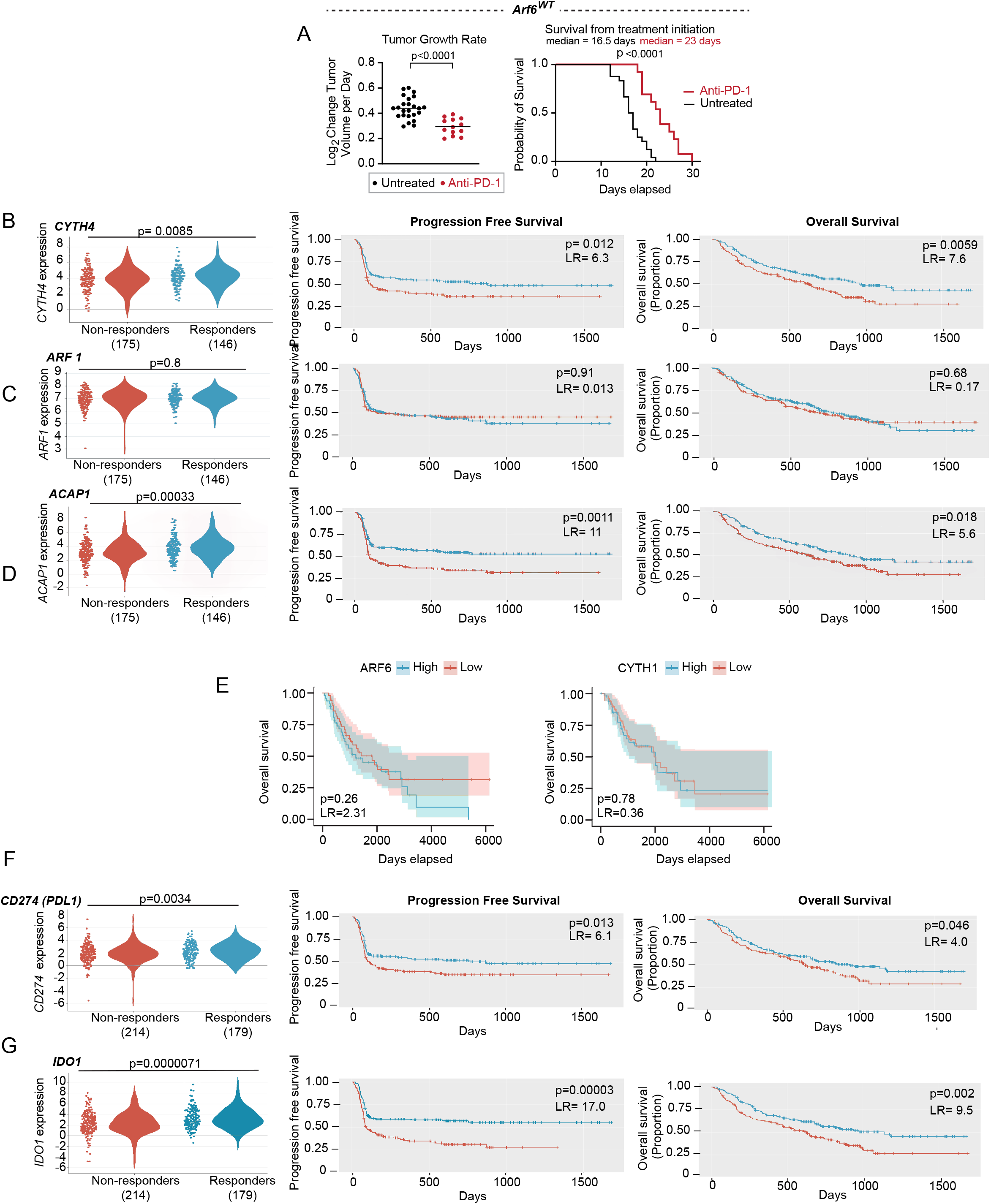
ICB treatment outcomes, related to Figure 6. **(A)** Systemic anti-PD-1 treatment initiated in *Arf6^WT^* mice with established tumors (up to 5mm in greatest dimension, 27-72mm^3^). Untreated controls (n=24), anti-PD-1 (n=13). Rate of tumor growth measured from initiation of treatment, Welch’s t-test. Survival (primary tumor reached 2cm) from initiation of treatment, Log-rank (Mantle-Cox) test. **(B-D)** Association of ICB treatment outcome in melanoma patients with mRNA levels of *CYTH4* **(B)**, *ARF1* **(C)** and *ACAP1* **(D)** in transcriptomes of pretreatment melanoma biopsies (CancerImmu expression analysis). **(E)** Lack of association of *ARF6* and *CYTH1* expression (tumor) with survival of non-ICB treated melanoma patients (TCGA, n=163). **(F-G)** Association of ICB treatment outcome in melanoma patients with mRNA levels of *CD274* **(F)**, *IDO1* **(G)** in transcriptomes of pretreatment melanoma biopsies, CancerImmu expression analysis, aggregated data from n=10 queried melanoma clinical studies, adjusted p-values, Benjamini and Hochberg procedure, LR= likelihood ratio (df=1).

